# Mitochondrial pyruvate import in astrocytes links anaplerosis to seizure resistance

**DOI:** 10.64898/2026.07.04.736458

**Authors:** Dario Garcia-Rodriguez, Sara Yunta-Sanchez, Marta Antequera-Düwel, Luisa Hidalgo-Lopez, Jesus Agulla, Leticia Sancha-Ortega, Rebeca Lapresa, Emilio Fernandez, Irene Martinez-Gallego, Adrian Sanchez-Gallego, Juan Fernandez-Garcia, Sandra Plaza-Garcia, Iiana Keren, Mohar Chattopadhyay, Simon Eaton, Simon J. R. Heales, Antonio Rodriguez-Moreno, Melanie Planque, Sarah-Maria Fendt, Pedro Ramos-Cabrer, Blanca I. Aldana, Angeles Almeida, Daniel Jimenez-Blasco, Juan P. Bolaños

## Abstract

Astrocytes are glycolytic cells that convert a substantial fraction of glucose-derived pyruvate into lactate, a metabolite implicated in supporting neuronal energy demand and modulating excitability, plasticity and memory. This view has placed astrocytic lactate production and export at the centre of astrocyte-neuron metabolic coupling, but whether mitochondrial pyruvate utilization in astrocytes is dispensable *in vivo* or fulfils an essential function in the intact brain remains unknown. Here we show that adult astrocyte-specific deletion of *Mpc2*, encoding an obligatory mitochondrial pyruvate carrier subunit, causes motor deficits, neuronal hyperexcitability and seizure-associated lethality. Metabolic profiling revealed pyruvate diversion toward alanine as an unsuccessful compensatory bypass, together with impaired tricarboxylic acid-cycle metabolism and an imbalance in neurotransmitter-related pools, including glutamate, glutamine and γ-aminobutyric acid. Thus, astrocytic mitochondrial pyruvate import is not primarily required for bioenergetic purposes but acts as a non-redundant anaplerotic gate that maintains neurotransmitter homeostasis, excitation-inhibition balance and seizure resistance *in vivo*.

## Main text

Astrocytes have long been recognized as metabolic partners of neurons. Whereas neurons are optimized for oxidative metabolism and actively suppress a strong glycolytic phenotype^1–3^, astrocytes retain high glycolytic capacity and respond to synaptic activity by increasing glucose uptake and lactate production^4–8^. Work over the last decade has reframed lactate as a physiological metabolite produced predominantly in astrocytes from glucose or glycogen in response to neuronal activity and transferred to neurons to satisfy energetic demand and modulate excitability, plasticity and memory^8–10^. We have recently shown that astrocytic glycolysis is required for cognition and that the glycolytic ATP-supported reverse-ATP synthase activity preserves mitochondrial membrane potential while limiting mitochondrial pyruvate decarboxylation and electron transport^11^. This observation raises the question as to whether the purpose of this metabolic feature of astrocytes is to drive mitochondrial pyruvate utilization to non-energetic alternative metabolic routes. The mitochondrial pyruvate carrier (MPC), formed by MPC1 and MPC2, is the canonical route for cytosolic pyruvate entry into the mitochondrial matrix^12,13^. Whilst in neurons this route feeds pyruvate dehydrogenase and oxidative ATP production^14^, in cultured astrocytes, mitochondrial pyruvate has a more specialized fate, namely pyruvate carboxylation to oxaloacetate, an anaplerotic reaction that replenishes tricarboxylic acid (TCA)-cycle intermediates to support *de novo* glutamate, glutamine and γ-aminobutyric acid (GABA)-related pools^15–17^. However, the physiological role of mitochondrial pyruvate uptake in astrocytes remains unexplored *in vivo*, leaving a fundamental gap in our understanding of how astrocytic pyruvate anaplerosis supports neurotransmitter balance, brain function and organismal health.

To address this question, we deleted *Mpc2* in adult astrocytes *in vivo* by delivering *PHP.eB-AAV-gfaABC1D-iCre-GFP* to *Mpc2^lox/lox^* mice, using the non-recombining vector as control (**Fig. 1a**). Immunomagnetic purification of astrocytes from the brain of the *Gfap-Mpc2-KO* (and wild type, WT) mice (**Fig. 1a; Extended Fig. 1a**) confirmed a marked reduction of MPC2 protein (**Fig. 1b**) and a concomitant decrease in pyruvate decarboxylation (**Fig. 1c**). However, neurons immunomagnetically purified from the *Gfap-Mpc2-KO* mice (**Extended Data Fig. 1a**) showed unaltered MPC2 protein (**Extended Data Fig. 1b**) and pyruvate decarboxylation (**Extended Data Fig. 1c**), indicating that this experimental approach disrupts mitochondrial pyruvate transport selectively in astrocytes. These *Gfap-Mpc2-KO* mice showed reduced exploration of the centre of an open field (**Fig. 1d**), impaired rotarod performance (**Fig. 1e**), without loss of strength (**Fig. 1f**) or cognitive impairment (**Fig, 1g,h**). However, these mice showed a loss of survival beginning around two months after viral transduction (**Fig. 1i**). The terminal phenotype was consistently preceded by seizure-like episodes (**Source Data Video**), and both decreased survival (**Extended Data Fig. 1d**) and motor dyscoordination (**Extended Data Fig. 1e,f**) were also observed in females. Immunocytochemical analysis of the hippocampus and cortex -two regions relevant for seizure activity-, revealed increased GFAP-positive area (**Fig. 1j**) consistent with reactive astrogliosis, whereas ß-TUBULIN III staining was unchanged (**Fig. 1j**) indicating preserved gross neuronal integrity. In line with these findings, brain from *Gfap-Mpc2-KO* mice showed reduced activity-dependent immediate-early genes *cFos* and *Arc mRNA* expression (**Extended Data Fig. 1g**), likely reflecting impaired neuronal transcriptional competence or metabolic exhaustion. Interestingly, CA1 pyramidal neurons from *Gfap-Mpc2-KO* mice fired action potentials at a lower threshold and generated more spikes in response to depolarizing current (**Fig. 1k; Extended Data Fig. 1h**). Together, these data indicate that MPC2 loss in astrocytes triggers a non-cell-autonomous state in which mitochondrial pyruvate-gating failure is accompanied by reactive astrogliosis, neuronal hyperexcitability and seizure-associated premature lethality.

**Fig. 1.**
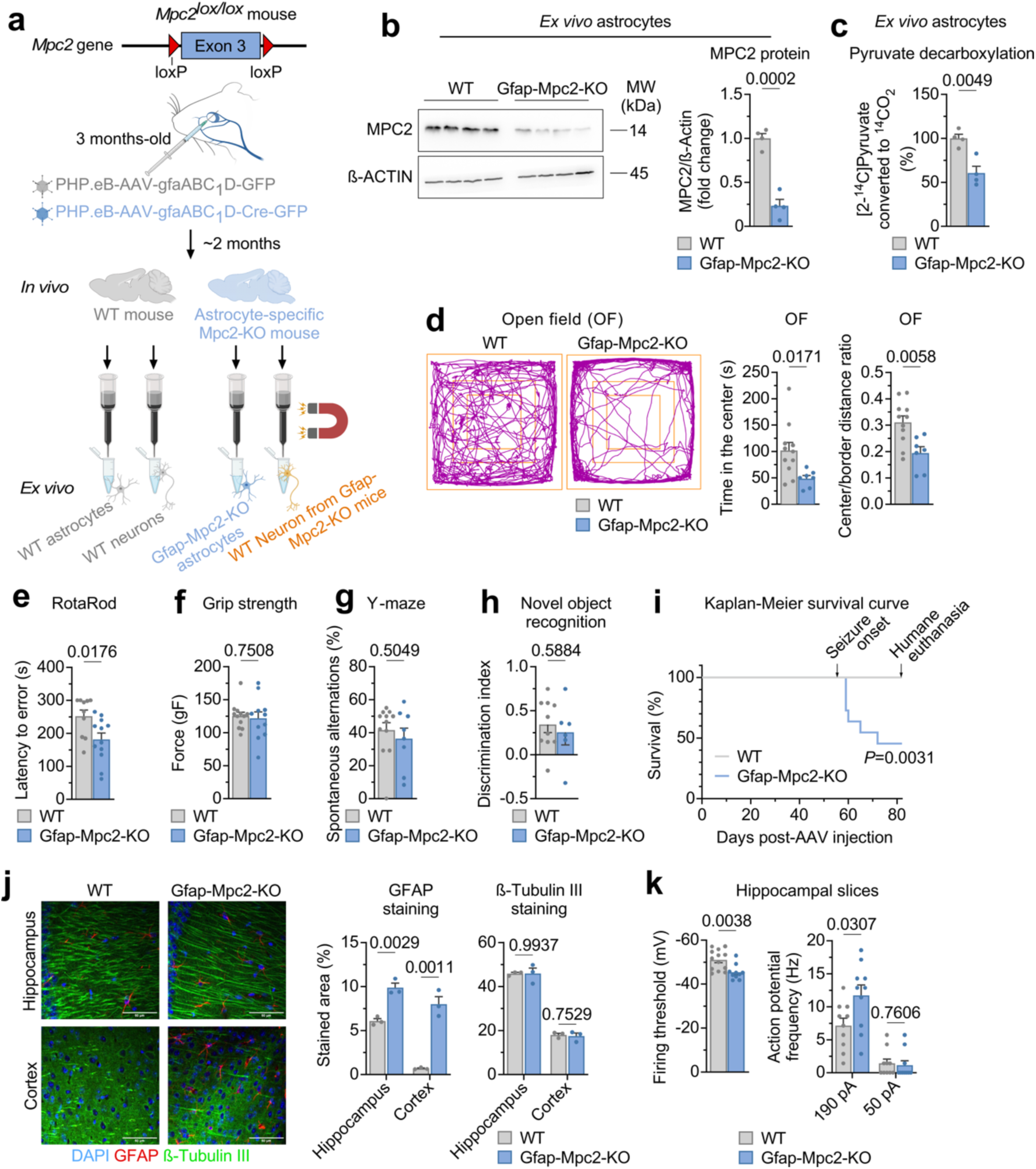
Astrocyte-specific *Mpc2* knockout (*Gfap-Mpc2-KO*) mice develops locomotion impairment and neuronal hyperexcitability-associated lethal seizures. **(a)** Strategy used to generate astrocyte-specific *Mpc2* knockout (*Gfap-Mpc2-KO*) mice and to immunomagnetically purify Mpc2-KO (and wild type, WT) astrocytes, and WT neurons from adult brain. Created with BioRender.com. **(b)** Western blot against MPC2 protein in astrocytes immunomagnetically purified from WT and *Gfap-Mpc2-KO* mouse brains; β-Actin was used as loading control (*left*). Quantification of MPC2 protein relative to β-Actin (*right*). Data are mean ± S.E.M. *P* value is indicated; n=4 mice per genotype; Unpaired Student’s *t*-test, two-tailed. **(c)** Pyruvate decarboxylation flux at the TCA cycle in astrocytes immunomagnetically purified from WT and *Gfap-Mpc2-KO*. Data are mean ± S.E.M. *P* value is indicated; n=4 mice per genotype; Unpaired Student’s *t*-test, two-tailed. Non-normalized values are 15.76 ± 0.76 nmol x h^−1^ x mg protein^−1^ for WT and 9.56 ± 1.29 nmol x h^−1^ x mg protein^−1^ for *Gfap-Mpc2-KO* mice.| **(d)** Open field test in WT and *Gfap-Mpc2-KO* male mice. Representative track plots for each genotype are shown (*left*). Quantification of time in the center and center/border distance ratio (*right*). Data are mean ± S.E.M. *P* values are indicated; n=11 (WT) or 7 (*Gfap-Mpc2-KO*) mice; Unpaired Student’s *t*-test, two-tailed. **(e)** Rotarod test in WT and *Gfap-Mpc2-KO* male mice. Data are mean ± S.E.M. *P* value is indicated; n=10 (WT) or 12 (*Gfap-Mpc2-KO*) mice; Unpaired Student’s *t*-test, two-tailed. **(f)** Grip strength in WT and *Gfap-Mpc2-KO* mice. Data are mean ± S.E.M. *P* value is indicated; n=12 (WT) or 11 (*Gfap-Mpc2-KO*) mice; Unpaired Student’s *t*-test, two-tailed. (gF, grams-Force). **(g)** Y-maze test in WT and *Gfap-Mpc2-KO* mice. Data are mean ± S.E.M. *P* value is indicated; n=12 (WT) or 8 (*Gfap-Mpc2-KO*) mice; Unpaired Student’s *t*-test, two-tailed. **(h)** Novel object recognition test in WT and *Gfap-Mpc2-KO* mice. Quantification of discrimination index is shown. Data are mean ± S.E.M. *P* value is indicated; n=10 (WT) or 6 (*Gfap-Mpc2-KO*) mice; Unpaired Student’s *t*-test, two-tailed. **(i)** Kaplan-Meier survival curve of WT and *Gfap-Mpc2-KO* male mice. *P* value is indicated; n=11 (WT) or 12 (*Gfap-Mpc2-KO*) mice; Logrank test. **(j)** Structural analysis of the cortex and hippocampus of WT and *Gfap-Mpc2-KO* mice. Representative staining of DAPI, GFAP and β-TUBULIN III is shown (*left*). Quantifications of the stained area of GFAP and β-TUBULIN III in cortex and hippocampus. Data are mean ± S.E.M. *P* values are indicated; n=3; Unpaired Student’s *t*-test, two-tailed. **(k)** Electrophysiological recordings in WT and *Gfap-Mpc2-KO* mice neurons from hippocampal slices. Quantifications of the firing threshold (*left*) and action potential frequency (*right*) are shown. Data are mean ± S.E.M. *P* values are indicated; n=13 (WT) or 10 (*Gfap-Mpc2-KO*) mice; Unpaired Student’s *t*-test, two-tailed.

Next, we assessed whether astrocytic MPC2 loss had any impact on the bioenergetics of neighboring neurons *in vivo*. Neurons immunomagnetically purified from *Gfap-Mpc2-KO* mice (**Fig. 1a; Extended Data Fig. 1a**) showed impaired basal and maximal mitochondrial respiration (**Fig. 2a**), indicating that astrocytic MPC2 loss non-cell-autonomously compromises neuronal bioenergetic fitness, likely through disrupted astrocyte-neuron metabolic coupling. To ascertain the metabolic consequences induced by MPC2 loss in astrocytes, we first performed spatial metabolomics in the hippocampus of *Gfap-Mpc2-KO* mice. The analysis revealed the accumulation of numerous metabolites (**Fig. 2b; Extended Data Fig. 2a**), including glycolytic intermediates such as fructose-1,6-bisphosphate (F-1,6-BP), phosphoenolpyruvate (PEP), and 2- or 3-phosphoglycerate (2-PG or 3-PG) (**Fig. 2b**), as well as fatty acids (**Extended Data Fig. 2a**). These findings are consistent with carbon accumulation upstream of mitochondrial pyruvate entry and diversion toward cytosolic metabolic fates. Serum lipoprotein, lipid, glycoprotein and low-molecular-weight metabolite profiles showed no broad systemic effect by genotype (**Extended Data Fig. 2b**), indicating that the metabolic basis of the lethal phenotype is largely confined to the brain. Analysis of common brain energy metabolites by *in vivo* 1H-MRS revealed increased alanine levels and a strong trend toward lactate accumulation -which was confirmed by GC/MS in brain homogenate (**Extended Data Fig. 2c**)-whereas glucose, glutathione, glutamine, glutamate and γ-aminobutyric acid (GABA) were reduced (**Fig. 2c**). These data further support carbon rerouting toward cytosolic metabolic fates and uncover a profound neurotransmitter imbalance in the *Gfap-Mpc2-KO* mouse brain. Notably, the marked increase in the glutamate/GABA ratio (**Fig. 2c, inset**) is consistent with a metabolic basis for the shift in excitation-inhibition balance toward the hyperexcitability that precedes lethal seizures. However, chronic intracerebroventricular administration of glutamine, aimed at restoring brain glutamate/GABA balance, failed to prevent seizures or mortality (**Fig. 2d**). Together, these data indicate that the brains of *Gfap-Mpc2-KO* mice undergo profound metabolic rewiring toward cytosolic pathways, accompanied by a marked neurotransmitter imbalance. Importantly, although this metabolic and neurochemical profile is consistent with neuronal hyperexcitability, seizures and lethality, the lack of rescue by local glutamine supplementation argues that the lethal phenotype cannot be explained by glutamine insufficiency alone, but instead reflects broader pathogenic consequences of impaired astrocytic mitochondrial pyruvate uptake.

**Fig. 2.**
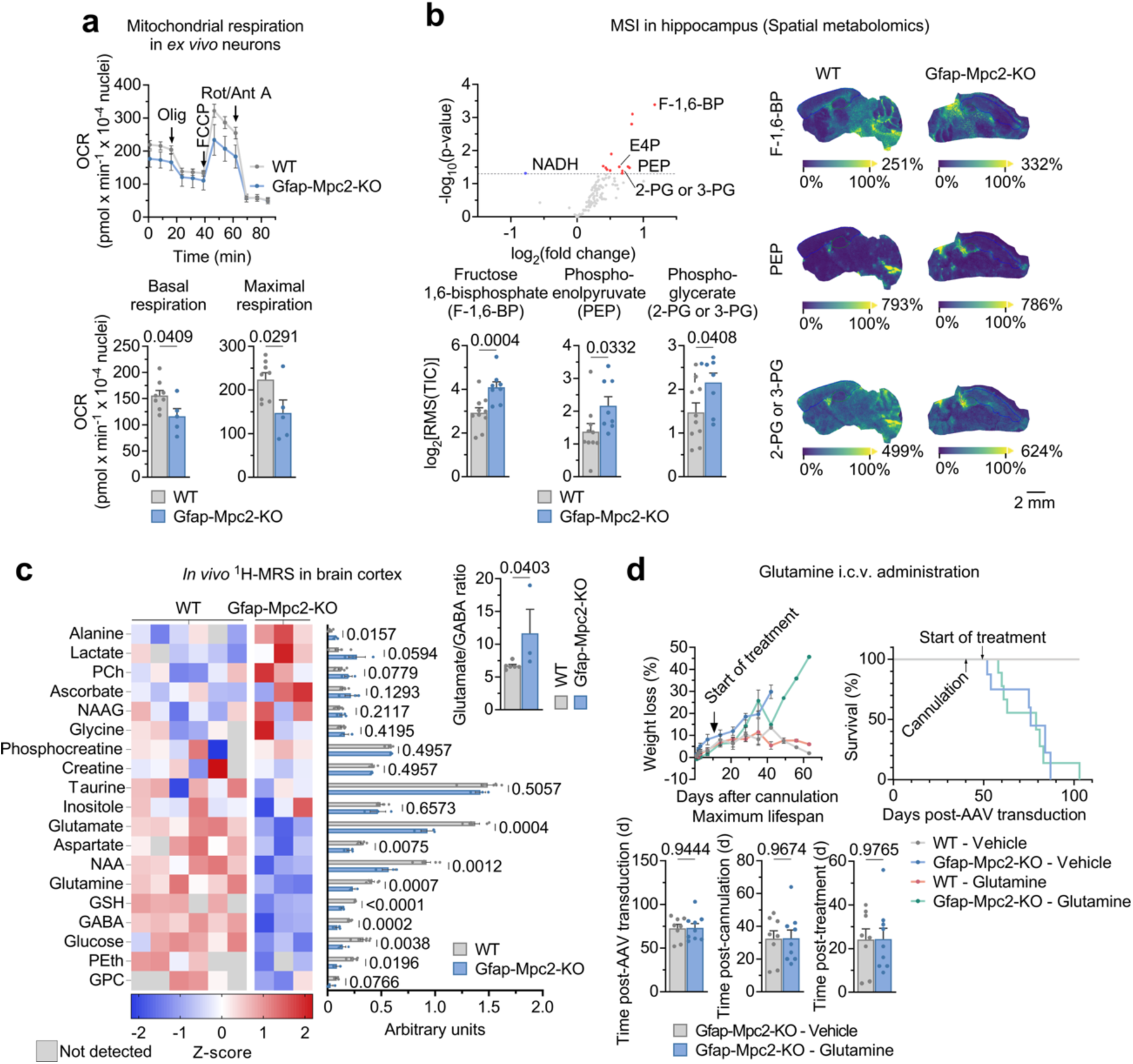
*Gfap-Mpc2-KO* mice displays brain metabolic rewiring including unbalanced excitatory/inhibitory neurotransmitter pool. **(a)** Oxygen consumption rate (OCR) analysis in neurons obtained from WT and *Gfap-Mpc2-KO* mice. *Top*: OCR traces; *bottom*: quantifications of basal and maximal respiration. Data are mean ± S.E.M. *P* values are indicated; n=8 (WT) or 5 (*Gfap-Mpc2-KO*) mice; Unpaired Student’s *t*-test, two-tailed. **(b)** Mass spectrometry imaging (MSI) in hippocampus of WT and *Gfap-Mpc2-KO* mice. *Top left:* volcano-plot for all the detected metabolites; *bottom left:* quantifications of the log2 of the Root Mean Squared (RMS) of the Total Ion Count (TIC) for the selected metabolites; *right*: representative scanned tissue slices from WT and *Gfap-Mpc2-KO* mice showing the intensity of the selected metabolites. Data are mean ± S.E.M. *P* values are indicated; n=10 (WT) or 8 (*Gfap-Mpc2-KO*) mice; Linear categorical model. **(c)** *In vivo* ^1^H-Magnetic resonance spectroscopy (MRS) in WT and *Gfap-Mpc2-KO* mice. *Left:* heatmap and quantifications of the detected metabolites through ^1^H-MRS. *Right, inset*: quantification of glutamate/GABA ratio levels. Data are mean ± S.E.M. *P* values are indicated; n=6 (WT) or 3 (*Gfap-Mpc2-KO*) mice; Unpaired Student’s *t*-test, one- or two-tailed. (PCh, phosphocholine; NAAG, N-acetyl-aspartyl-glutamate; NAA, N-acetyl-aspartate; GSH, glutathione; GABA, γ-aminobutyric acid; PEth, phosphotethanolamine, GPC, glycerophosphocholine). **(d)** Intracerebroventricular (I.c.v) injections of glutamine in WT and *Gfap-Mpc2-KO* mice. *Top left*: body weight loss of WT and *Gfap-Mpc2-KO* mice treated with vehicle or glutamine after cannulation; *top right*: Kaplan-Meier survival curve of WT and *Gfap-Mpc2-KO* mice treated with vehicle or glutamine; *bottom*: quantification of maximum lifespan of WT and *Gfap-Mpc2-KO* mice treated with vehicle or glutamine. Data are mean ± S.E.M. *P* values are indicated; n=4 (WT – Vehicle and WT - Glutamine), 8 (*Gfap-Mpc2-KO* - Vehicle) or 9 (*Gfap-Mpc2-KO* - Glutamine) mice; Unpaired Student’s *t*-test, two-tailed.

To assess the metabolic alterations occurring specifically in astrocytes, we next generated primary astrocytes cultures from *Mpc2^lox/lox^* mice and transduced them with an adenovirus expressing Cre recombinase or an empty vector as control (**Fig. 3a**). This approach successfully induced loss of MPC2, together with the concomitant loss of MPC1 (**Fig. 3b; Extended Data Fig. 3a**), consistent with the interdependence of both carrier subunits^18,19^. RNA-sequencing analysis of these cells confirmed reduced *Mpc2* and *Mpc1 mRNA* levels, together with *Gfap mRNA* upregulation, in *Mpc2-KO* astrocytes (**Fig. 3c**). Notably, this analysis also revealed increased expression of *Slc38a1 mRNA* (**Fig. 3c**), encoding the glutamine uptake transporter SNAT1 (sodium-coupled neutral amino acid transporter 1), which is predominantly restricted to neurons under physiological conditions^20^, as well as *Slc1a3 mRNA*, encoding the astrocyte-enriched glutamate–aspartate transporter GLAST, also known as excitatory amino acid transporter 1 (EAAT1) in humans (**Fig. 3c**). Together with the marked reduction in glutamine, glutamate and aspartate levels observed in the brain of *Gfap-Mpc2-KO mice* (**Fig. 2c**), these transcriptional adaptations suggest that loss of mitochondrial pyruvate carrier drives astrocytes to increase the uptake and utilization of these amino acids. To directly test this possibility in cultured astrocytes, we performed [U-^13^C]glucose tracing. This analysis revealed reduced incorporation of glucose-derived carbon into the TCA-cycle intermediates citrate, α-ketoglutarate, malate and succinate, as well as into the amino acids glutamate and aspartate, in *Mpc2-KO* astrocytes (**Fig. 3d**), strongly supporting impaired anaplerotic biosynthesis. To determine whether the reduction in brain glutamine levels reflected increased utilization or reduced synthesis, we next assessed glutamine-dependent mitochondrial respiration, which was unchanged in *Mpc2-KO* astrocytes (**Extended Data Fig. 3b**). Likewise, the rates of [1-^14^C]glutamine oxidation to ^14^CO2 and [1-^13^C]glutamate oxidation to ^13^CO2 were not altered by MPC2 loss (**Extended Data Fig. 3c,d**). Together, these data indicate that the reduced brain levels of glutamine, glutamate and aspartate primarily result from impaired biosynthesis rather than increased consumption, helping to explain why *in vivo* glutamine administration was insufficient to prevent lethal seizures (**Fig. 2d**).

**Fig. 3.**
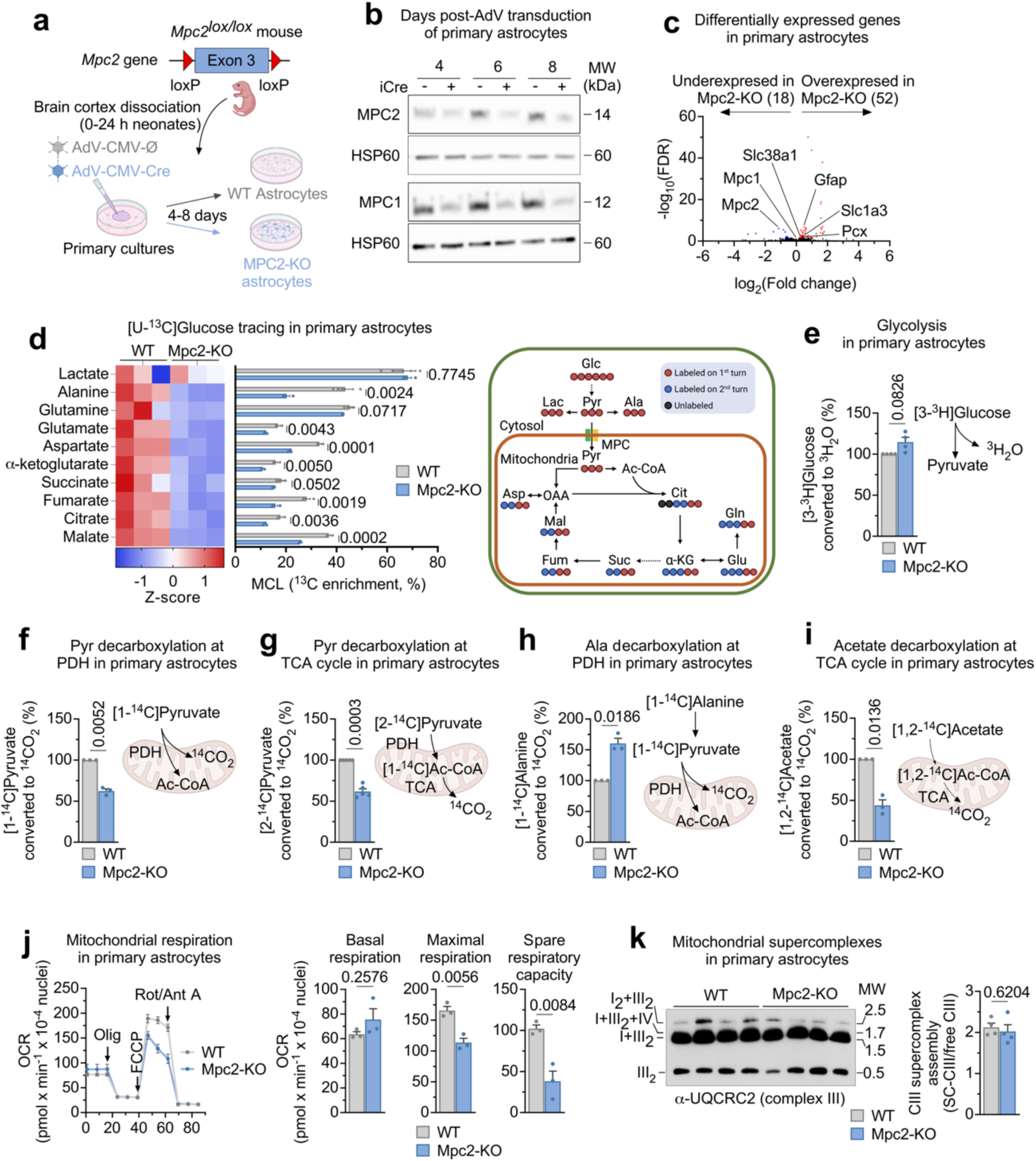
Lack of *Mpc2* in primary astrocytes impairs anaplerotic TCA cycle without bioenergetic deficiency. **(a)** Strategy used to obtain *Mpc2* knockout astrocytes in primary culture. Created with BioRender.com. **(b)** Representative Western blot showing MPC2 and MPC1 protein levels 4, 6 and 8 days after AdV-CMV-Cre transduction in immunomagnetically purified mitochondria from primary astrocytes; HSP60 was used as loading control. **(c)** RNA-Sequencing in WT and *Mpc2-KO* primary astrocytes. Selected genes are shown. n=3 biologically independent samples per genotype; Wald test with local fitting. **(d)** [U-^13^C]Glucose tracing in WT and *Mpc2-KO* primary astrocytes. *Left*: heatmap and *right*: molecular carbon labeling (MCL) quantification of the detected metabolites. Data are mean ± S.E.M. *P* values are indicated; n=3 biologically independent samples per genotype; Unpaired Student’s *t*-test, two-tailed. **(e)** Glycolytic flux in WT and *Mpc2-KO* primary astrocytes. Data are mean ± S.E.M. *P* value is indicated; n=4 biologically independent samples per genotype; Unpaired Welch’s *t*-test, two-tailed. Non-normalized values are 384.76 ± 118.84 nmol x h^−1^ x mg protein^−1^ for WT and 445.49 ± 140.60 nmol x h^−1^ x mg protein^−1^ for *Mpc2-KO* astrocytes. **(f)** Pyruvate decarboxylation flux at the PDH in WT and *Mpc2-KO* primary astrocytes. Data are mean ± S.E.M. *P* value is indicated; n=3 biologically independent samples per genotype; Unpaired Welch’s *t*-test, two-tailed. Non-normalized values are 146.09 ± 57.26 nmol x h^−1^ x mg protein^−1^ for WT and 93.56 ± 37.46 nmol x h^−1^ x mg protein^−1^ for *Mpc2-KO* astrocytes. **(g)** Pyruvate decarboxylation flux at the TCA cycle in WT and *Mpc2-KO* primary astrocytes. Data are mean ± S.E.M. *P* value is indicated; n=5 biologically independent samples per genotype; Unpaired Welch’s *t*-test, two-tailed. Non-normalized values are 26.52 ± 5.65 nmol x h^−1^ x mg protein^−1^ for WT and 16.05 ± 3.05 nmol x h^−1^ x mg protein^−^1 for *Mpc2-KO* astrocytes. **(h)** Alanine decarboxylation flux at the PDH in WT and *Mpc2-KO* primary astrocytes. Data are mean ± S.E.M. *P* value is indicated; n=3 biologically independent samples per genotype; Unpaired Welch’s *t*-test, two-tailed. Non-normalized values are 11.51 ± 2.18 nmol x h^−1^ x mg protein^−1^ for WT and 18.33 ± 3.12 nmol x h^−1^ x mg protein^−1^ for *Mpc2-KO* astrocytes. **(i)** Acetate decarboxylation flux at the TCA cycle in WT and *Mpc2-KO* primary astrocytes. Data are mean ± S.E.M. *P* value is indicated; n=3 biologically independent samples per genotype; Unpaired Welch’s *t*-test, two-tailed. Non-normalized values are 5.79 ± 1.90 nmol x h^−1^ x mg protein^−1^ for WT and 2.37 ± 0.51 nmol x h^−1^ x mg protein^−1^ for *Mpc2-KO* astrocytes. **(j)** Oxygen consumption rate (OCR) analysis in WT and *Mpc2-KO* primary astrocytes. *Left*: OCR traces; *right*: quantifications of basal respiration, maximal respiration and spare respiratory capacity. Data are mean ± S.E.M. *P* values are indicated; n=3 biologically independent samples per genotype; Unpaired Student’s *t*-test, two-tailed. **(k)** Analysis of the mitochondrial respiratory chain complex III superassembly. *Left*: Blue-native gel electrophoresis images showing complex III (CIII) containing supercomplexes (SC-CIII) and free CIII in WT and *Mpc2-KO* astrocytes. *Right*: Quantification of SC versus free CIII. Data are mean ± S.E.M. *P* value is indicated; n=4 biologically independent samples per genotype; Unpaired Student’s *t*-test, two-tailed.

To further understand the metabolic rewiring occurring in *Mpc2-KO* astrocytes, we next assessed the glycolytic flux using [3-^3^H]glucose -a *bona fide* proxy for this pathway^2^. This analysis revealed a slight, albeit not statistically significant, increased glycolytic flux (**Fig. 3e**). In contrast, [1-^14^C]pyruvate decarboxylation was reduced (**Fig. 3f**), without changes in phosphorylated pyruvate dehydrogenase (pPDH) levels (**Extended Data Fig. 3e**), indicating impaired mitochondrial pyruvate entry and oxidation in the absence of altered PDH inhibitory phosphorylation. Consistently, ^14^CO2 production from [2-14C]pyruvate, which reflects pyruvate oxidation through the TCA cycle, was similarly decreased (**Fig. 3g**). Intriguingly, pyruvate-dependent respiration under basal conditions was unaltered (**Extended Data Fig. 3f**). However, MPC2-deficient astrocytes appeared to partially bypass mitochondrial pyruvate entry by increasing the oxidation of [1-14C]alanine (**Fig. 3h**), consistent with cytosolic pyruvate transamination to alanine, mitochondrial alanine uptake, and subsequent transamination back to pyruvate followed by PDH-dependent decarboxylation. Nevertheless, this compensatory route was insufficient to restore TCA-cycle flux, as shown by reduced ^14^CO2 production from [1,2-14C]acetate (**Fig. 3i**), further supporting impaired mitochondrial carbon metabolism in *Mpc2-KO* astrocytes. Notably, *mRNA* levels of the anaplerotic enzyme pyruvate carboxylase (*Pcx*) were increased in *Mpc2-KO* astrocytes (**Fig. 3c**), suggesting an unsuccessful compensatory response to defective pyruvate-dependent anaplerosis. In line with these findings, basal mitochondrial respiration (**Fig. 3j**), mitochondrial membrane potential (ΔΨm) (**Extended Data Fig. 3g**), mitochondrial ROS (mROS) production (**Extended Data Fig. 3h**), the NAD^+^/NADH(H^+^) ratio (**Extended Data Fig. 3i**) and respiratory-chain supercomplex assembly (**Fig. 3k**) were largely preserved. However, maximal respiration and spare respiratory capacity were impaired (**Fig. 3j**), suggesting that mitochondrial pyruvate oxidation in astrocytes becomes particularly important under increased energetic demand. Consistent with this, the contribution of succinate as an oxidative substrate was increased twofold under uncoupling conditions (**Extended Data Fig. 3j**), indicating a compensatory reliance on alternative mitochondrial substrates. Thus, astrocytes lacking MPC2 preserve respiratory organization and basal mitochondrial function, but fail to sustain full respiratory flexibility and glucose-derived pyruvate-dependent anaplerotic metabolism.

Although astrocytes adapt to the loss of mitochondrial pyruvate uptake by rewiring anaplerotic metabolism, this adaptation may non-cell-autonomously compromise neighboring neurons. Accordingly, wild type neurons cocultured with *Mpc2-KO* astrocytes (**Fig. 4a**) showed reduced levels of the antioxidant glutathione (GSH) (**Fig. 4b**), likely reflecting limited glutamine availability (**Fig. 2c**) for *de novo* neuronal GSH biosynthesis^21,22^. Concomitantly, these neurons displayed increased mROS and reduced ΔΨm, without detectable loss of neuronal viability (**Fig. 4b-d**). These findings recapitulate key features observed in the brains of *Gfap-Mpc2-KO* mice (**Fig. 1j,k**; **Fig. 2a**) and suggest that the anaplerotic impairment caused by loss of astrocytic mitochondrial pyruvate metabolism becomes particularly detrimental under the metabolic demands imposed by the close astrocyte-neuron interactions. Interestingly, astrocyte-specific MPC2 loss led to the accumulation of a broad lipid signature in the brain, as revealed by spatial metabolomics (**Extended Data Fig. 2a**). We therefore asked whether fatty acid utilization was impaired in *Mpc2-KO* astrocytes. Basal fatty acid-dependent respiration was unchanged (**Extended Data Fig. 4a; Fig. 4f, left panel, inset**). In contrast, under the bioenergetic stress imposed by mitochondrial uncoupling, the contribution of fatty acids to maximal respiration and spare respiratory capacity was markedly increased (**Fig. 4f, two right panels, insets**), indicating that fatty acids are recruited as alternative oxidative substrates when respiratory demand rises. Consistent with preserved basal fatty acid oxidation, both fatty acid β-oxidation and β-oxidation-coupled acetyl-CoA decarboxylation fluxes were unchanged in *Mpc2-KO* astrocytes (**Fig. 4g,h**). However, fatty acid conversion into ketone bodies was reduced (**Fig. 4i**), suggesting that MPC2 loss selectively limits the ability of astrocytes to channel fatty acid-derived carbon into ketogenesis. To determine whether systemic ketone body availability could mitigate the *in vivo* phenotype, we chronically administered a ketogenic diet. Despite efficient engagement of ketogenic metabolism, as shown by body weight gain and blood ketone and glucose levels (**Extended Data Fig. 4b–d**), ketogenic diet did not alter the increased lactate (**Extended Data Fig. 4e**) or the decreased *Fos* and *Arc mRNA* (**Extended Data Fig. 4f**) brain levels, failing to preserve motor coordination or prevent disease progression in *Gfap-Mpc2-KO* mice (**Fig. 4j**). Together with the lack of rescue by glutamine supplementation, these findings indicate that neither fatty acid-derived acetyl-CoA, systemic ketone-body availability, lactate nor glutamine supply can substitute for the continuous astrocytic anaplerotic conversion of pyruvate into oxaloacetate required to sustain the endogenous glutamate-glutamine-GABA axis (**Extended data Fig. 4g**). Thus, mitochondrial pyruvate uptake is not merely one of several interchangeable carbon inputs in astrocytes, but a non-redundant anaplerotic route that preserves neuronal metabolic homeostasis and organismal viability *in vivo*.

**Fig. 4.**
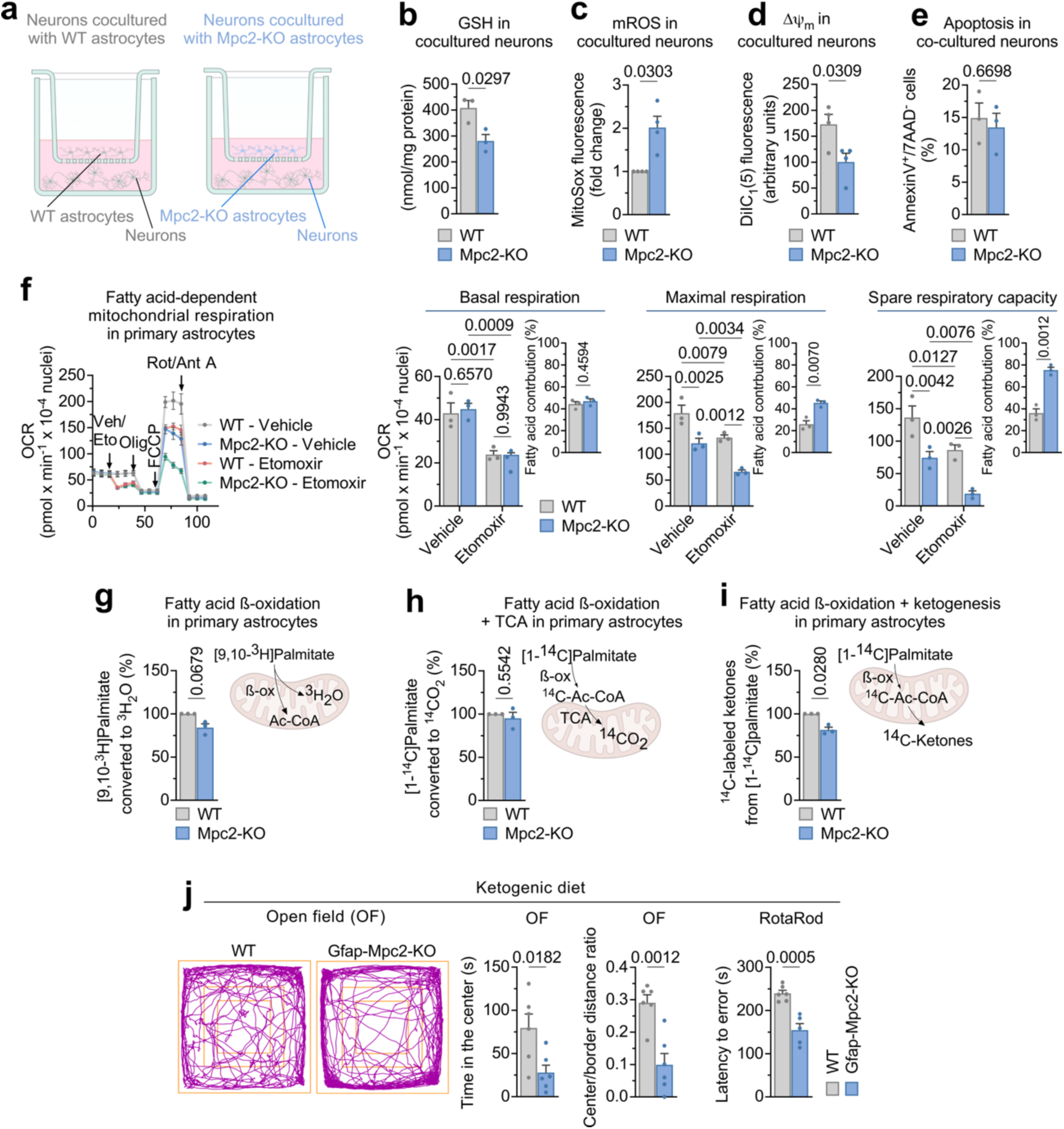
Astrocyte-specific *Mpc2* loss causes neuronal redox and bioenergetic stress *via* fatty acids-derived ketones-independent mechanism. **(a)** Strategy used to obtain primary neurons co-cultured with WT and *Mpc2* knockout astrocytes. Created with BioRender.com. **(b)** Glutathione (GSH) levels in neurons co-cultured with WT and *Mpc2-KO* astrocytes. Data are mean ± S.E.M. *P* value is indicated; n=3 biologically independent samples per genotype; Unpaired Student’s *t*-test, two-tailed. **(c)** Mitochondrial ROS (mROS) levels in neurons co-cultured with WT and *Mpc2-KO* astrocytes. Data are mean ± S.E.M. *P* value is indicated; n=4 biologically independent samples per genotype; Unpaired Welch’s *t*-test, two-tailed. **(d)** Mitochondrial membrane potential (ΔΨm) levels in neurons co-cultured with WT and *Mpc2-KO* astrocytes. Data are mean ± S.E.M. *P* value is indicated; n=4 biologically independent samples per genotype; Unpaired Student’s *t*-test, two-tailed. **(e)** Apoptosis in neurons co-cultured with WT and *Mpc2-KO* astrocytes. Data are mean ± S.E.M. *P* value is indicated; n=3 biologically independent samples per genotype; Unpaired Student’s *t*-test, two-tailed. **(f)** Fatty acid-dependent mitochondrial respiration in WT and *Mpc2-KO* primary astrocytes. *Left*: OCR traces; *right*: quantification of basal respiration, maximal respiration and spare respiratory capacity; *right, insets*: fatty acid contribution to each metabolic parameter. Data are mean ± S.E.M. *P* values are indicated; n=3 biologically independent samples per genotype; Two-way ANOVA followed by Tukey or Unpaired Student’s *t*-test, two-tailed. **(g)** Fatty acid β-oxidation flux in WT and *Mpc2-KO* primary astrocytes. Data are mean ± S.E.M. *P* value is indicated; n=3 biologically independent samples per genotype; Unpaired Welch’s *t*-test, two-tailed. Non-normalized values are 6.99 ± 2.64 nmol x h^−1^ x mg protein^−1^ for WT and 5.94 ± 2.32 nmol x h^−1^ x mg protein^−1^ for *Mpc2-KO* astrocytes. **(h)** Fatty acid β-oxidation and TCA cycle decarboxylation flux in WT and *Mpc2-KO* primary astrocytes. Data are mean ± S.E.M. *P* value is indicated; n=3 biologically independent samples per genotype; Unpaired Welch’s *t*-test, two-tailed. Non-normalized values are 1.14 ± 0.26 nmol x h^−1^ x mg protein^−1^ for WT and 1.12 ± 0.33 nmol x h^−1^ x mg protein^−1^ for *Mpc2-KO* astrocytes. **(i)** Ketogenesis flux in WT and *Mpc2-KO* primary astrocytes. Data are mean ± S.E.M. *P* value is indicated; n=3 biologically independent samples per genotype; Unpaired Welch’s *t*-test, two-tailed. Non-normalized values are 0.34 ± 0.03 nmol x h^−1^ x mg protein^−1^ for WT and 0.28 ± 0.02 nmol x h^−1^ x mg protein^−1^ for *Mpc2-KO* astrocytes. **(j)** Open field and RotaRod tests in WT and *Gfap-Mpc2-KO* mice treated with a ketogenic diet. *Left:* representative track plots for each genotype. *Center*: quantifications of the time spent in the center and the center/border distance ratio. *Right:* RotaRod test quantification. Data are mean ± S.E.M. *P* values are indicated; n=6 mice per genotype; Unpaired Student’s *t*-test, two-tailed.

In conclusion, our findings show that astrocytes do not require mitochondrial pyruvate primarily as a dominant oxidative ATP source, but as a strategically positioned carbon input that sustains pyruvate carboxylase-dependent anaplerosis^16^. We further reveal that in the absence of MPC *in vivo*, astrocytes preserve glycolysis, lactate and alanine production, basal mitochondrial organization and respiratory integrity, and can recruit fatty-acid oxidation under energetic stress. However, these compensatory routes fail to replace the continuous conversion of pyruvate into oxaloacetate required to maintain the endogenous glutamate-glutamine-GABA axis. Consequently, neurotransmitter precursor homeostasis collapses, GABA falls disproportionately *in vivo*, excitation-inhibition balance shifts toward hyperexcitability, and mice develop lethal seizures. Thus, astrocytic mitochondrial pyruvate uptake reveals an anaplerotic requirement that is largely masked in resting cultured cells but becomes indispensable in the intact brain, where astrocytes must continuously sustain neurotransmitter precursor renewal imposed by active astrocyte-neuron metabolic coupling.

This conclusion extends the astrocyte-neuron metabolic coupling framework, including the astrocyte-neuron lactate shuttle and its broader role in brain function^23–25^. Astrocytic pyruvate metabolism therefore contains two complementary branches, namely, an exportable lactate branch that supports neuronal activity, memory and behaviour, and an internal MPC-dependent anaplerotic branch that preserves transmitter-pool stability. This distinction also helps position astrocytic pyruvate gating within seizure metabolism. Astrocytes are increasingly recognized as causal regulators of seizure susceptibility through potassium buffering, water flux, glutamate uptake, adenosine metabolism, gap junction coupling and intermediary metabolism^26,27^, whereas ketogenic and medium-chain triglyceride strategies demonstrate that fuel selection can modulate neuronal excitability through mechanisms involving mitochondrial biogenesis, antioxidant responses, AMPA receptor inhibition, astrocyte lactate/ketone shuttling and astrocyte-derived glutamine support for neuronal GABA synthesis^20,28–30^. Our data help explain why such interventions may fail when the missing function is not fuel availability *per se*, but pyruvate-derived oxaloacetate production. Thus, acetyl-CoA-rich substrates, ketone bodies or glutamine supplementation cannot substitute for astrocytic pyruvate-dependent anaplerosis, because none of these interventions restores the *de novo* oxaloacetate-generating step required to replenish neurotransmitter precursor pools in the brain^15–17^. Together with recent evidence that neuronal pyruvate oxidation can tune circuit performance through long-term memories-dependent mitochondrial Ca²⁺ handling^14^, our study argues that pyruvate gates have cell-type-specific meanings in the brain. Neurons use pyruvate oxidation to enhance activity-dependent energy production^31^, whereas astrocytes use mitochondrial pyruvate entry to preserve the anaplerotic economy that prevents activity from becoming pathological. By placing astrocytic MPC at the interface between pyruvate metabolism, neurotransmitter homeostasis, excitation-inhibition balance and organismal viability, this work identifies anaplerosis -not ATP production alone- as a central determinant of astrocyte-dependent network stability and a potential metabolic entry point for future therapeutic strategies in neurological disease.

## Methods

### Mpc2^lox/lox^ mice

All protocols were performed according to the European Union Directive 86/609/EEC and Recommendation 2007/526/EC, regarding the protection of animals used for experimental and other scientific purposes, enforced in Spanish legislation under the law 6/2013. Protocols were approved by the Bioethics Committee of the University of Salamanca in accordance with the Spanish legislation (RD53/2013). All mice used in this study were of the C57Bl/6J background. *Mpc2^lox/lox^*mice were obtained from The Jackson Laboratories (strain #032118). Animals were bred at the Animal Experimentation Facility of the University of Salamanca in cages (maximum of six animals per cage) with a 12 h light-dark cycle (light from 08:00 h). The humidity was 45–65%, and the temperature was 20-25 °C. Animals were fed *ad libitum* with a standard solid diet (Envigo-Harlan Tekland Global 18% Protein Rodent Diet, USA; 18% proteins, 3% lipids, 58.7% carbohydrates, 4.3% cellulose, 4% minerals, and 12% humidity) or a ketogenic diet (D10070801, Research diets, USA; 90% fat, 10% proteins) and free access to water.

### Blood glucose and ketone bodies determination

Glucose and ketone bodies concentration in blood from WT and Gfap-Mpc2-KO mice were determined with a Freestyle Optimus Neo glucometer (Abbott) coupled to glucose measuring strips (252346.3, FreeStyle) and ketone bodies measuring strips (252353.1, FreeStyle).

### *In vivo* generation of astrocyte-specific *Mpc2* knockout mice

This was carried out using a validated adeno-associated virus (AAV) strategy^32^. Essentially, AAV particles of the PHP.eB capsid (serotype), known to efficiently transduce the central nervous system via intravenous injection^33^, expressing Cre recombinase driven by the astrocyte-specific short glial-fibrillary acidic protein (GFAP) promoter (PHP.eB-AAV-gfaABC1D-Cre-GFP) were administered intravenously (50 µl aliquots of a phosphate-buffered saline solution containing 0.001% Pluronic® F-68, Sigma-Aldrich, and 1 × 10^11^ viral genomes, VG) through the retro-orbital sinus to 3 months-old *Mpc2^lox/lox^* mice under a brief sevoflurane anesthesia (Sevorane, AbbVie, Spain, at 6% for initiation followed by ∼3% for maintenance in air with supplement O2 and NO2 -0.4 and 0.8 l/min, respectively- using a gas distribution column, Hersill H-3, Spain, and a vaporizer, InterMed Penlons Sigma Delta, UK). We used the retro-orbital sinus intravenous route because of the higher success rate observed when compared with the tail or temporal ones^34^. Siblings of wild-type (WT) mice received equivalent amounts of the same AAV particles that did not harbor Cre recombinase. AAV were acquired from the Viral Vector Facility of the University of Zurich (v95 for AAV-GfaABC1D-GFP and v232 for AAV-GfaABC1D-Cre-GFP). Mice were used 2 months after AAV injections. The efficacy of this approach at promoting recombination exclusively in astrocytes was previously confirmed in our laboratory using *Cpt1a^lox/lox^*mice^35^.

### Mouse cannulation and glutamine treatment

To infuse glutamine directly into the brain ventricles, a cannula was inserted to WT and Gfap-Mpc2-KO mice 1.5 months after AAV injection. Briefly, animals were anesthetized with 4% sevoflurane in a mixture of O2:NO2 (30:70%) using a gas distribution column. Skin from the top of the head was cut and the skull was perforated with a drill coupled to a stereotaxic instrument. The cannula was placed by mounting 26-gauge short single guide cannula (C315GS-5/SPC, Bilaney) in the stereotaxic instrument, and it was inserted at 0.22 mm anterior to bregma, 1.0 mm lateral to midline, and 2.5 mm ventral to dura. Cannula was fixed with dental cement and glue, and a single dummy cannula (C315DCS-5/SPC, Bilaney) was placed as a cap. The cut skin was sutured, and mice were individualized in cages after the surgery. Glutamine infusions started 10 days after the surgery. To perform the glutamine infusion, brain ventricular volume was estimated to be 8 µl, and glutamine was prepared to reach a final concentration in the cerebrospinal fluid of 2 mM. Glutamine (G5763, Sigma) was diluted in 0.9% NaCl and loaded into 10 µl Hamilton syringes. Four Hamilton syringes were placed on a CMA 4004 microdialysis syringe pump (CMA Microdyalisis, Kista, Sweden), that infused 2 µl of volume at a flow of 1 µl/min. Hamilton syringes were coupled to a cannula tubing (C315CT/PKG, Bilaney) and this in turn is coupled to a 33-gauge short single internal cannula (C315IS-5/SPC, Bilaney). To couple the internal cannula to the guide cannula, mice were anesthetized with sevoflurane anesthesia at 4% in a mixture of O2:NO2 (30:70%).

### Primary cultures of astrocytes

Astrocytes in primary culture were obtained from the cortex of 0-24 h old *Mpc2^lox/lox^* mouse neonates^36^. Cell suspension was seeded in 175 cm^2^ plastic flasks in low glucose (5.5 mM) Dulbecco’s Modified Eagle’s Medium (DMEM) supplemented with 10% fetal bovine serum and 4 mM glutamine and incubated at 37 °C in a humidified 5% CO2-containing atmosphere. To detach non-astrocytic cells, after 7 days in vitro (DIV), the flasks were shaken at 150 r.p.m. overnight. The supernatant was discarded, and the attached, astrocyte-enriched cells were reseeded at 0.5-1 × 10^5^ cells/cm^2^ in the appropriate plates. Cells were used at 16 DIV. This method has previously reported a ∼100% purity of astrocytes in the culture^35^.

### Neuron-astrocyte coculture

Individual primary cultures of mouse cortical neurons were prepared from E14.5 day-old c57BL6/J mice^36^, seeded at 2.0 × 10^5^ cells per cm^2^ in six-well plates coated with poly-D-lysine (10 μg/ml) and incubated in Neurobasal A (A2477501, Thermo Fisher Scientific) supplemented with 2 mM glutamine, 5.5 mM glucose, 0.22 mM pyruvate and 2% B27 supplement. Cells were incubated at 37 °C in a humidified 5% CO2-containing atmosphere. 72 h after plating, the medium was replaced. To obtain neuron-astrocytes cocultures, transwell membrane inserts (4.5 cm^2^, 0.4 µm pore size; Corning) containing WT or Mpc2-KO astrocytes at 5 days post adenoviral transduction were placed over the neuronal culture. Neuron-astrocyte coculture was left untreated for 3 days, and neurons were used on day 6^35^.

### Generation of *Mpc2* knockout astrocytes in primary culture

This was carried out by transducing 9 DIV primary astrocytes, obtained from *Mpc2^lox/lox^* mice, with adenoviral particles harboring Cre recombinase driven by the ubiquitous cytomegalovirus (CMV) promoter (AdV-CMV-Cre). Astrocytes from the same cultures transduced with equivalent amounts of the same AdV lacking Cre recombinase (AdV-CMV-Ø) were used as WT astrocytes. The AdV were purchased from the viral repository of the University of Iowa (VVC-U of Iowa-5 for AdV-CMV-Cre; VVC-U of Iowa-272 for AdV-CMV-Ø). Cells were used 8 days after transduction.

### Genotyping by polymerase chain reaction (PCR)

For *Mpc2^lox/lox^* genotyping, a PCR with the following primers was performed; primer 1, 5’-AGGCACCAAGGAAGATATGGT-3’ (forward) and primer 2, 5’-TTAAGTACCTGGAAAGTTCACTCAG-3’ (reverse), resulting in a 360 bp band for *Mpc2^lox/lox^* mice and 277 bp for wild type. PCR conditions were 3 min at 94°C, 38 cycles of 15 s at 94°C, 20 s at 60°C, 30 s at 72°C, and final extension of 10 s at 72°C. Products were resolved in 3% agarose gel using the 100 bp DNA ladder plus (Thermo Fisher Scientific).

### Immunomagnetic purification of astrocytes and neurons from adult brain

Mouse adult brain (minus cerebellum and olfactory bulb) was dissociated using the adult mouse brain dissociation kit (Miltenyi Biotec). The tissue, once clean, was fragmented with a sterile scalpel in 2 ml per hemisphere of a disintegration solution (Earle’s Balanced Salt Solution, EBSS, 116 mM NaCl, 5.4 mM KCl, 1.5 mM MgSO4, NaHCO3 26 mM, NaH2PO4 1.01 mM, glucose 4 mM, phenol red 10 mg/l, pH 7.2, supplemented with albumin 14.4 μl/ml and DNase type I 26 μl/ml and trypsin 10.8μl/ml), and it was trypsinized at 37°C in a thermostated bath for 5 minutes, shaking frequently to avoid decantation of the tissue. It was further mechanically disintegrated by trituration using a 5 ml serological pipette for 5 times. Then, the suspension was returned to the thermostated bath for 10 minutes, shaking frequently. Trypsin activity was stopped by adding 10% fetal serum, before centrifuging the tissue at 700 g for 5 minutes in a microfuge at 4 °C. Once the enzymatically disintegrated tissue had been decanted, the pellet was resuspended in a trypsin-free disintegration solution (EBSS + 13 μl/ml DNase + 20 μl/ml albumin) for mechanical trituration using a Pasteur pipette. 5 passages were performed per volume of 4 ml and per hemisphere. The supernatant was centrifuged for 3 min at 700 g. Once a homogeneous suspension of individualized adult neural cells was achieved, cell population separations were performed using MACS® Technology using either the astrocyte-specific anti-ACSA-2 Microbead Kit or the neuron-specific Neuron Isolation Kit, according to manufacturer’s protocol (MACS® technology). It has been previously confirmed the identity of the isolated fractions by Western blotting against astrocytic (GFAP), neuronal (MAP2)-specific markers, and the purity with microglial (Iba1) and oligodendroglial (OLIG2)-specific markers^35^.

### Immunomagnetic purification of mitochondria from astrocytic cultures

Immunomagnetic mitochondrial isolation was performed using the mitochondria isolation kit, mouse tissue (130-096-946, Miltenyi Biotec). The protocol was adapted to be applied to cultured astrocytes. Astrocytes cultured on 145 cm^2^ dish plates were collected by trypsinization. Cells were resuspended in 500 μl of Buffer A (sucrose 83 mM, MOPS 10 mM; pH 7.2) supplemented with the protease inhibitory cocktail (aprotinin (A1153, Sigma) 50 μg/ml; leupeptin (L2884, Sigma) 50 μg/ml; trypsin inhibitor from soy (T9128, Sigma) 50 μg/ml; Nα-tosyl-L-lysine chloromethyl ketone (TLCK; T7254, Sigma) 100 μM; phenylmethanesulfonyl fluoride (PMSF; P7626, Sigma) 100 μM; N-p-tosyl-L-phenylalanine chloromethyl ketone (TPCK; T4376, Sigma) 100 μM; *o*-phenanthroline (P9375, Sigma) 1 mM and pepstatin A (P4265, Sigma) 50 μg/ml ) and phosphatase inhibitor cocktail (Phosphatase inhibitor cocktail 3 (P0044, Sigma)), and grinded with an electric homogenizer coupled to a glass-teflon Potter-Elvehjem. Samples were placed on a glass mortar and grinded performing 15 rod passes. Buffer B (sucrose 250 mM, MOPS 30 mM; pH 7.2) supplemented with the protease and a phosphatase inhibitor cocktails was added to the homogenate, and it was subsequently diluted in 10 ml of Separation Buffer 1X. 50 μl of α-TOMM22 antibody bound to magnetic microbeads were added. Homogenized samples were incubated with the microbeads for 1 hour at 4 °C in an orbital shaker. Labeled samples were loaded onto LS columns (130-042-401, Miltenyi Biotec) placed in a QuadroMACS Separation Unit (130-090-976, Miltenyi Biotec). Afterwards, columns were washed 3 times with 3 ml of Separation Buffer 1X. Lastly, columns were removed from the separator and placed on 2 ml Eppendorf tubes.

1.5 ml of Separation Buffer 1X was added and immediately pushed through the column with the help of a plunger to elute labeled mitochondria. Samples were centrifuged at 13,000 g 2 minutes at 4 °C, supernatant was discarded, and the pellet was resuspended in 1 ml of Storage Buffer. Samples were again centrifuged at 13,000 g 2 minutes at 4 °C, and supernatant was discarded to obtain purified mitochondria.

### Determination of metabolic fluxes

To assess metabolic fluxes, we used radiometric approaches. To do this, astrocytes were seeded in 8 cm^2^ flasks (353108, Fisher) hanging a microcentrifuge tube containing either 1 ml benzethonium hydroxide (Sigma) (for ^14^CO2 equilibration) or 1 ml H2O (for ^3^H2O equilibration). All incubations were carried out in in Krebs-Ringer phosphate buffer (KRPG; NaCl 145 mM, Na2HPO4 5.7 mM, KCl 4.86 mM, CaCl2 0.54 mM, MgSO4 1.22 mM, glucose 5.5 mM; pH 7.35) at 37 °C in the air-thermostatized chamber of an orbital shaker. Pyruvate oxidation was assessed by measuring the rate of production of ^14^CO2 from [1-^14^C]pyruvate and [2-^14^C]pyruvate using 0.25 μCi/ml of either of the radioisotopic tracers in KRPG supplemented with 1 mM pyruvate for 90 minutes. Alanine oxidation was assessed by measuring the rate of production of ^14^CO2 from [1-^14^C]alanine using 0.5 μCi/ml of [1-^14^C]alanine in KRPG supplemented with 1 mM alanine for 90 minutes. The glycolytic flux was measured by assaying the rate of ^3^H2O production from [3-^3^H]glucose using 2 μCi/ml of D-[3-^3^H]glucose for 3 h^2^. To determine TCA cycle activity, the rate of production of ^14^CO2 from [1,2-^14^C]acetate was measured using 0.5 μCi/ml of [1,2-^14^C]acetate in KRPG supplemented with 0.5 mM sodium acetate for 90 min. The rate of fatty acids ß-oxidation -i.e., the conversion of fatty acids to acetyl-Coenzyme A- was performed by analyzing the rate of ^3^H2O production from [9,10-^3^H]palmitate (1 μCi/ml) in KRPG supplemented with 30 µM palmitate for 3 h^37–39^. To measure the carbon flux from fatty acids to CO2, - which jointly assesses ß-oxidation and acetyl-CoA decarboxylation in the TCA cycle-, cells were incubated in KRPG supplemented with 30 µM palmitate and 0.25 µCi/ml of [1-^14^C]palmitic acid^40 68^. The rate of ^14^CO2 production from [1-^14^C]glutamine was assessed using [1-^14^C]glutamine (0.5 μCi/ml) in KRPG supplemented with 2 mM glutamine. All the incubations were terminated with 0.2 ml 20 % perchloric acid. Cells were further incubated for 60 min to allow benzethonium hydroxide to trap ^14^CO2 or 72 h to allow for ^3^H2O equilibration with H2O present in the central microcentrifuge tube. The ketone bodies were extracted as a non-volatile, acid-soluble product from [1-14C]palmitic acid incubations. Thus, after stopping the reaction with perchloric acid, 1 ml of medium was collected and transferred to a delipidated tube containing 8 volumes of chloroform/methanol (2:1, v/v) and 2 volumes of KCl (0.1 M). After shaking, it was centrifuged for 5 min at 3000 g, and 2 ml of the upper aqueous phase were transferred to a new tube with 4 volumes of the previous mixture, to obtain a purer sample. It was centrifuged again, and 2 1 ml aliquots of the aqueous phase were taken and transferred to vials with scintillation fluid for radioactivity determination (ketogenesis). The ^14^CO2, ^3^H2O and ^14^C-labeled ketone bodies were then measured by liquid scintillation counting Tri-Carb 4810 TR (PerkinElmer). In all cases, the specific radioactivity was used for the calculations. Under these experimental conditions, 70% of the produced ^14^CO2 and 28% of the produced ^3^H2O were recovered and were considered for the calculations^2^.

### Determination of [1-^13^C]Glutamate conversion to ^13^CO2

To assess [1-^13^C]Glutamate conversion to ^13^CO2, a DMEM formulation was prepared consisting of DMEM powder (D5030, Sigma) 8.3 g/l, glucose 5 mM, HEPES 15 mM, NaHCO3 2.9 mM, phenol red (P5530, Sigma) 21.5 μM; pH 7.4. Astrocytes cultured cells in 6 well plates were washed once with PBS and incubated overnight with DMEM-HEPES supplemented with 10 % FBS. Afterwards, cells were incubated with DMEM-HEPES supplemented with [1-13C]Glutamate (604968, Sigma) 1 mM and sealed with a 3 ml layer of heavy mineral oil (330760, Sigma Aldrich) to prevent ^13^CO2 loss. Astrocytes were incubated for 6 hours, and 100 μl media aliquots were taken at 1-, 3- and 6-hours incubation time by inserting a Hamilton syringe through the mineral oil. Samples were collected in rubber-sealed *Exetainer*™ vials (539W, Labco Ltd). Samples were thawed at room temperature, and 100 μL 1M hydrochloric acid was injected through the septum into each vial to release CO2 from the medium. Vials were centrifuged for 30 s at 500 g. Samples were then analyzed on a GasBench II coupled to a Thermo Delta-XP isotope-ratio mass spectrometer (Thermo-Finnigan, Bremen, Germany). Ten repeat injections were carried out per sample, with ^13^CO2/^12^CO2 ratios measured against Vienna Pee Dee Belemnite (VPDB) using a calibrated CO2 reference gas. Following this, ^13^CO2/^12^CO2 ratios were then converted to mole percent excess using absolute molar ratio of ^13^C to ^12^C (0.0111796) in VPDB. The change in mole percent excess was then converted to pmol CO2 generated using the volume of medium and concentration of bicarbonate (2.9 mM) present^41^.

### Brain lactate levels

Hemispheres were stored at -80°C until analysis. Mouse brains were weighed and homogenized 1:10 (wt/vol) in phosphate-buffered saline, 1 mmol/l EDTA and 5 mmol/l Tris, pH 7.4 using a glass homogenizer. Brain homogenate (50 µL) were added to 225 µL acetonitrile, which was vortexed and centrifuged for 5 minutes at 15,000 *g*. The supernatant was dried under N2. Samples were resuspended in 200 µL 100% ethanol and redried. Samples were derivatised in ethyl acetate (30µL) and *N,O*-bis(trimethylsilyl)-trifluoroacetamide (BSTFA)/10% trimethylchlorosilane (TMCS) (30 µL) at 37°C for 30 minutes. Lactate was analysed by GC/MS (Thermo ISQ7610 with Trace 1310 GC, RXI-5Sil column (30m x 0.25mm I.D, 0.25mM film thickness), inlet 250°C, helium flow 1.2 ml/min, oven temperature 60°C for 1 minute, to 140°C at 10°C/min, then to 240°C at 40°C/min. Samples were analysed in electron impact mode, with selected ion monitoring at m/z 219.

### Mitochondrial ROS

Mitochondrial ROS were determined with the fluorescent probe MitoSox^TM^ Red (M36008, Thermo Fisher Scientific). Cultured cells were incubated with 2 μM of MitoSox for 30 min at 37 °C in a 5% CO2 atmosphere in KRPG. The cells were then washed with phosphate-buffered saline (PBS; 136 mM NaCl; 2.7 mM KCl; 7.8 mM Na2HPO4·2H2O; 1.7 mM KH2PO4; pH 7.4) and collected by trypsinization. MitoSox fluorescence intensity was assessed by flow cytometry (FACScalibur, FL-3 channel, flow cytometer, CellQuestTM software, BD Biosciences) and values were normalized by the mitochondrial membrane potential measured, since it is known that the MitoSox probe is affected by the ΔΨm42.

### Flow cytometric analysis of cell death

Primary neurons cocultured with either WT or Mpc2-KO astrocytes were incubated with APC-conjugated annexin V and 7-aminoactinomycin D (7-AAD) (Becton Dickinson Biosciences) to quantitatively determine the percentage of apoptotic and necrotic cells by flow cytometry. Cell suspensions were stained with annexin V–APC and 7-AAD in binding buffer (100 mM HEPES, 140 mM NaCl, 2.5 mM CaCl2), according to the manufacturer’s instructions, and 10^4^ cells were analyzed, in three replicates per condition, on a FACSCalibur flow cytometer (15-mW argon ion laser, CellQuest software, Becton Dickinson Biosciences), using FL4 and FL3 channels, respectively. Annexin^+^/7AAD^−^ cells were considered to be apoptotic. Data were expressed as percentages.

### NAD^+^ and NADH(H^+^) determinations

To determine NAD^+^/NADH(H^+^) levels, a commercial kit (MAK460, Sigma) was used following the manufacturer’s instructions. Excitation intensities of 530 nm and emission at 585 nm were determined. The 96-well plate was incubated for 10 minutes in the dark at RT, and the measurement was carried out at time 0 and after 10 minutes. To calculate the NAD^+^ or NADH(H^+^) concentration, a standard line was run in parallel from 0 (blank) to 1 μM NAD^+^ or NADH(H^+^) from which the sample concentrations were extrapolated.

### Determination of glutathione

Cells were lysed with 1% (wt/vol) sulfosalicylic acid (S0640, Sigma) and centrifuged at 13,000 *g* for 5 min at 4 °C, and the supernatants were used for determining glutathione by using GSSG (0–50 µM) as a standard. Total glutathione was measured in reaction buffer [0.1 mM Na2HPO4, 1 mM EDTA, 0.3 mM DTNB (D8130, Sigma), 0.4 mM NADPH (N1630, Sigma) and glutathione reductase (G3664, Sigma) at 1 U/ml, pH 7.5] by recording the increase in absorbance at 405 nm after the reaction of GSH with DTNB for 15 min at 15 s intervals using a Varioskan Flash reader (Thermo Fisher Scientific).

### *In vitro* RNA extraction

RNA extraction was carried out using the commercial *Genelute Mammalian Total RNA Kit* (RTN350, Sigma Aldrich) according to the manufacturer’s instructions. Briefly, cells were lysed and scraped with a mixture of Lysis Solution:β-mercaptoethanol (Sigma) 100:1 (vol/vol). Subsequently, an equivalent volume of ethanol 70 % in diethylpyrocarbonate (DEPC) water, RNase free, was added. The mixture was then passed through a *Genelute Binding Column*, where the RNA was adhered. The column was washed once with *Wash Solution 1*. Once the RNA was separated, the column was treated with 100 U of DNase I (04716728001, Roche) resuspended in the *DNAse I buffer* for 15 minutes at RT. After incubation, to remove all DNA residues, columns were washed three times, once with *Wash Solution 1* and twice with *Wash Solution 2 Concentrate/ethanol*, allowing for complete column drying. Finally, the column content was eluted with 50 μl of nuclease-free H2O (BP2484, Fisher Scientific) by centrifugation at 16,000 *g* for 1 minute. Purified RNA concentration was measured using a *UV-Vis Nanodrop 2000* spectrophotometer (Thermo Fischer Scientific).

### RNA-Sequencing

RNA samples from astrocytes were sent to *Macrogen Inc.* to perform RNA-Sequencing analysis. The library used for RNA sequencing was *TruSeq Stranded Total RNA* with *Ribo-Zero for Mouse Samples* on the platform *NovaSeq 6000* (Illumina). The free online software from the Universidad de Salamanca RaNA-Seq^43^ was used to perform sequencing alignment, transcript annotation and gene quantification. Prior to the analysis, quality control for each set of reads and a principal component analysis were used to discard samples that did not group appropriately with their experimental condition.

### qPCR with reverse transcription

RNA from mice hemispheres was extracted using the RNA TRI reagent according to the manufacturer’s instructions. Briefly, the brain tissue was resuspended and homogenized in 1.7 ml of TRI reagent (93289, Sigma Aldrich). Lysates were then centrifuged at 12,000 *g* for 10 minutes at 4 °C and the supernatant was collected and incubated for 15 min at RT with 170 μl of 1-bromo-3-chloropropane (B9673, Sigma Aldrich) After a 10 min 12,000 *g* centrifugation at 4 °C, the aqueous phase was collected in a new tube and 1 ml of 2-propanol (109634, Sigma Aldrich) was added. Samples were left to precipitate for 10 minutes at RT and then centrifuged at 12,000 *g* for 10 minutes at 4 °C. Supernatant was discarded, and pellets were rinsed with 1.7 ml of ethanol 75 %. Pellets were subsequently resuspended in nuclease-free H2O and incubated at 55 °C for 13 minutes. RNA concentration and its purity was measured using a *UV-Vis Nanodrop 2000* spectrophotometer. To evaluate the expression of the target genes, the commercial *Power SYBR® Green RNA-to-CT™ 1-Step Kit* (4389986, Applied Biosystems) was used. 100 ng of RNA were loaded in a final reaction volume mix of 20 μl, which consisted of 10 μl of *SYBR Green*, 0.16 μl of RT Enzyme, 0.6 μl of oligonucleotides 10 μM, and nuclease-free H2O. All reactions were performed in triplicate using the *Mastercycler ep Realplex thermocycler* (Eppendorf). The primers used were (forward and reverse, respectively) 5′-GGGAATGGTGAAGACCGTGT-3′ and 5′-CCGCATGGACTGTGGTCATGA-3′ for *cFos;* 5′-CACTCTCCCGTGAAGCCATT-3′ and 5-TCCTCCTCAGCGTCCACA-TA-3′ for *Arc,* and 5′-CAAGATCATTGCTCCTCCTG-3′ and 5′-CTGCTTGCTGATCCACATCT-3′ for *β-actin*. The relative mRNA levels were calculated using the ΔΔ*C*t method, and the resulting normalized values were expressed as the fold change *versus* WT.

### Oxygen consumption rate assessment

Oxygen consumption rates (OCR) were measured in real-time in an XFe24 Extracellular Flux Analyzer (Seahorse Bioscience; *Seahorse Wave Desktop software* 2.6.1.56). This equipment measures the extracellular medium O2 flux changes of cells seeded in XFe24-well plates. Regular cell medium was removed and washed once with DMEM running medium (DMEM supplemented with 10 mM glucose, 2 mM L-glutamine, 1 mM sodium pyruvate, 5 mM HEPES, pH 7.4). Primary astrocytes were incubated at 37°C without CO2 for 1 hour to allow cells to pre-equilibrate with the assay medium, while *ex vivo* isolated neurons from adult brain were immediately analyzed after being seeded. For the fuel capacity test, UK-5099 (PZ0160, Sigma) 10 µM, BPTES (SML0601, Sigma), etomoxir E1905, Sigma) 100 µM rotenone (R8875, Sigma) 1 µM and antimycin A (A8674, Sigma) 2.5 µM diluted in DMEM running medium were used. Fuel capacity was calculated as indicated by the manufacturer. For the MitoStress test, oligomycin (O4876, Sigma) 2.5 µM, FCCP (C2920, Sigma) 4 µM and a mixture of rotenone 1 µM and antimycin A 2.5 µM, diluted in DMEM running medium were loaded into port-A, port-B, and port-C, respectively. In some cases, etomoxir 100 µM and dimethyl malonate (DMM; 136441, Sigma) 10 mM were loaded into port-A, shifting the oligomycin, FCCP and rotenone/antimycin mix to ports-B, -C and -D respectively. The sequence of measurements was as follows. Basal level of oxygen consumption rate (OCR) was measured 3 times, and then port-A was injected and mixed for 3 min, after OCR was measured 3 times for 3 min. Same protocol with port-B, port-C and port-D. OCR was measured after each injection to determine mitochondrial or non-mitochondrial contribution to OCR. After the experiment, the number of nuclei in each well was determined using an Operetta CLS high content screening microscope (Perkin Elmer) to normalize the OCR measurements. Each sample was measured in 3–4 replicas. Experiments were repeated 3–4 times in biologically independent culture preparations and 5–8 independent neuron isolations for *ex vivo* neuronal OCR measurements. Basal respiration was determined by the OCR rate before the first injection minus OCR after rotenone/antimycin injection. Maximal respiration was determined by the OCR rate after FCCP injection minus OCR after rotenone/antimycin injection. Spare respiratory capacity was calculated as the maximal OCR minus the basal OCR. Fatty acids and succinate contribution to each respiratory parameter was calculated as: (non-inhibited OCR-inhibited OCR)/(non-inhibited OCR) and expressed as a percentage.

### Protein determinations

Protein samples were quantified by the BCA protein assay kit (Thermo Fisher Scientific) using BSA as a standard.

### Western Blotting

Samples were lysed in RIPA buffer (1% sodium dodecylsulfate, 10 mM ethylenediaminetetraacetic acid (EDTA), 1 % (vol/vol) Triton X-100, 150 mM NaCl and 10 mM Na2HPO4, pH 7.0), supplemented with protease and phosphatase inhibitor cocktails. Aliquots of sample lysates (20-80 μg of protein) were subjected to SDS/PAGE on an acrylamide gel (MiniProtean; Bio-Rad), including *PageRuler Prestained Protein Ladder* (Thermo). The resolved proteins were transferred electrophoretically to nitrocellulose membranes (0.2 µm, BioRad) at 90 V for 120 minutes. Membranes were blocked with 5% (wt/vol) low-fat milk (Sveltesse, Nestle) in TTBS (20 mM Tris, 150 mM NaCl, and 0.1% (vol/vol) Tween 20, pH 7.5) for 1 h. After blocking, membranes were immunoblotted with primary antibodies overnight at 4 °C. After incubation with horseradish peroxidase-conjugated goat anti-rabbit IgG (1/20,000, sc-2020, Santa Cruz Biotechnologies), goat anti-mouse IgG (1/30,000, 1858413, Pierce), and rabbit anti-goat IgG (1/20,000, sc-2701, Santa Cruz Biotechnologies), membranes were immediately incubated with the enhanced chemiluminescence kit *WesternBright ECL* (Advansta), or *SuperSignal West Femto* (Thermo) before exposure to *Fusion FX Vilber transilluminator*. At least three biologically independent replicates were always performed, although only one representative Western blot is shown in the main figures. The protein abundances of all Western blots per condition were measured by densitometry of the bands on the films using *ImageJ 1.54 software* (National Institutes of Health). They were normalized by loading control protein. The resulting values were used for statistical analysis. Uncropped scans of Western blots replicas are shown in the *Source Data file*.

### Primary antibodies for Western blotting

Immunoblotting was performed with anti-MPC1 (1/500) (14462; Cell Signaling), anti-MPC2 (1/500) (46141; Cell Signaling), anti-PDHA1 (1/1,000) (3205; Cell Signaling), anti-p-PDHA1(Ser293) (1/1,000) (31866; Cell Signaling), anti-GFAP (1/500) (G6171; Sigma), anti-β-TUBULIN III (1/500) (ab18207, Abcam), anti-MAP2 (1/500) (ab11268, Abcam), anti-HSP60 (1/1,000) (ab46798; Abcam), anti-β-actin (1/30,000) (A5441; Sigma) and anti-GAPDH (1/20,000) (AM4300; Ambion).

### Mitochondrial isolation

To obtain the mitochondrial fraction for blue native gel electrophoresis, cell pellets were frozen at −80 °C and homogenized (ten-twelve strokes) in a glass-Teflon Potter–Elvehjem homogenizer in buffer A. The same volume of buffer B was added to the sample, and the homogenate was centrifuged (1,000 *g*, 5 min) to remove unbroken cells and nuclei. Centrifugation of the supernatant was then performed (12,000 *g*, 3 min) to obtain the mitochondrial fraction, which was washed in buffer C (320 mM sucrose; 1 mM EDTA and 10 mM Tris-HCl; pH 7.4)^44^. Mitochondria were suspended in buffer D (1 M 6-aminohexanoic acid and 50 mM Bis-Tris-HCl, pH 7.0) for blue native gel electrophoresis.

### Blue native gel electrophoresis

For the assessment of supercomplex I-III organization, digitonin-solubilized (4 g/g) mitochondria (10–20 μg) were loaded in NativePAGE Novex 3–12% (vol/vol) gels (Life Technologies). The electrophoresis was carried out at 40 V overnight. Next, a direct electrotransfer was performed, followed by immunoblotting against mitochondrial the complex III antibody UQCRC2 (1/1,000) (ab14745; Abcam). Direct transfer of BNGE was performed after soaking the gels for 20 min (4 °C) in carbonate buffer (10 mM NaHCO3; 3 mM Na2CO3·10H2O; pH 9.5–10). Proteins were transferred to polyvinylidene fluoride (PVDF) or 0.2 mm nitrocellulose membranes (Hybond^®^, Amersham Biosciences) and were carried out at 300 mA, 60 V, 1.5 h at 4 °C in carbonate buffer.

### Mouse perfusion, immunohistochemistry, and image analysis

Animals were deeply anesthetized by intraperitoneal injection of a mixture (1:3) of xylazine hydrochloride (Rompum, Bayer) and ketamine hydrochloride/chlorbutol (Imalgene; Merial) using 1 ml of the mixture per kg of body weight, and then perfused intraaortically with 0.9 % NaCl followed by 5 mL/g body weight of fixative solution [4 % (wt/vol) paraformaldehyde, depolymerized with NaOH and stabilized with 10 % (v/v) methanol, in 0.1 M phosphate buffer, pH 7.4]. After perfusion, brains were dissected out sagittally into two parts and postfixed, using fixative solution, for 24 h at 4 °C. Afterwards, brains were cryoprotected in 30 % (wt/vol) sucrose in PBS. After cryoprotection, 10, 20, and 40-μm-thick sagittal sections were obtained with a cryostat (Leica; CM1950 AgProtect). Sections were then incubated with primary antibodies anti-GFAP (1/500) (G6171; Sigma) and β-TUBULIN III (1/500) (Ab18207, Abcam), in in 0.2 % Triton X-100 (Sigma) and 10 % goat serum (Jackson ImmunoReseach) in 0.1 M PBS for 72 h at 4°C. Later, fluorophore-conjugated secondary antibodies goat anti-rabbit-Cy2 (1/500) (111-225-144, Jackson lmmunoresearch), goat anti-mouse-Cy3 (1/500) (115-165-003, Jackson lmmunoresearch) in 0.05 % Triton X-100 and 2 % goat serum in 0.1 M PB for 2 h at room temperature^45^. For nuclear staining, sections were incubated with DAPI 0.1 mg/ml in PB for 10 min to stain nuclei. Finally, sections were mounted with Fluoromont (Sigma) aqueous mounting medium. Confocal images were taken with a laser scanning confocal microscope (*Olympus IX81 Confocal spinning disk*, Roper Scientific) with three lasers of 405, 491, and 561 nm and equipped with a digital camera (Evolve). All images acquired by confocal microscopy were processed and, after deconvolution, analyzed using the free software FIJI (ImageJ64). The quantification was performed on Z-Project micrographs of three different regions of the CA1 from each hippocampus and from the brain cortex, taken from three serial sagittal sections per mouse. Both the percentage of area occupied by β-TUBULIN III and GFAP were measured.

### Behavioral tests

Male and female mice (2 months post AAV injections, i.e., 5 months old) were left to acclimatize in the room for not less than 30 minutes at the same time slot of the day (9 am-14 pm). Both sexes were evaluated separately at first. After determining there was no sexual dimorphism in the knocked-out animal behavior, males and females were both evaluated indistinctively. Tracking was carried out once at a time and carefully cleaning the apparatus with 70% ethanol between trials to remove any odor cues. An ANY-box^®^ core (ANY-Maze, Stoelting Europe) was used, which contained a light grey base and an adjustable perpendicular stick holding a camera and an infrared photo-beam array connected to an AMi-maze^®^ interface to track the animal movement and to detect rearing behavior, respectively. For the *Openfield* test, a 40 cm x 40 cm x 35 cm (w, d, h) black infrared transparent Perspex insert was used, and the arena was divided into two zones, namely border (8 cm wide from the border of the arena) and center (the remaining area). The test lasted for 10 min, and the distance travelled and the time spent in each zone were measured. The *Rotarod test* (Rotarod apparatus, Model 47600, Ugo Basile) was used to analyze motor balance and coordination. Mice were previously trained for two consecutive days, and after they were subjected to the test. The rotarod conditions were a gradual acceleration from 4 to 40 r.p.m., reaching the final speed at 270 s. The day of the test, mice latency to fall from the rotarod or to skip three turns by grappling the rotarod was recorded. Mice grip strength was analyzed using a grip strength meter^46^ (Bioseb, Vitrolles, France). This meter consists of a force detector coupled to a metal rack. Animals gripped with the front paws to the rack, and a gentle push from the tail was performed to assess the grip strength to the metal rack. Results are expressed in grams-Force (gF) To analyze the short-term memory, we used the *Novel Object Recognition test* (Stoelting). Mice were accustomed to the ANY-box^®^ core for 10 min, 24 hours before the test. Mice were left to explore two identical equidistant objects (familiar object) for 5 min (the familiarization phase) and returned for 30 min into its cage. One object was replaced by another of similar size (novel object), and mice were returned to the arena to explore the objects for another 5 min (the test phase). The software scored the investigation of the objects when the head of the animal was looking at the object within a distance of 20 mm. The ability to recognize the novel object was determined as discrimination index (DI) calculated as [DI = (TN – TF) / (TN + TF)], where TN is the time spent exploring the new object and TF is the time spent exploring the familiar object. For the *Y-maze test* (working memory test), we used a maze composed of three arms in the form of a Y shape labeled A, B, and C, respectively. Each arm is 30 cm long, 5 cm wide, and 12 cm high. Animals were placed at the end of arm A and allowed to move freely through the maze for 5 minutes^6^. The ANY-maze software considered an arm entry was counted when 90% of the mouse body was inside the arm. Correct spontaneous alternation was defined as the number of sequential entry triads in three different arms (ABC, ACB, BAC, BCA, CAB, CBA). The percentage of correct spontaneous alternations was calculated as follows: [SA = (TS/(TAE-2)*100], where TS is the total number of correct spontaneous alternations and TAE is the total number of arm entries. The number of evaluated animals per test is specified in the figure legends.

### Electrophysiology recordings

Hippocampal slices were prepared as previously described^47–49^. Mice were anesthetized with isoflurane 2 % and decapitated for slice preparation. Briefly, after decapitation, the whole brain containing the two hippocampi was removed into ice-cold solution (I) consisting of NaCl 126 mM, KCl 3 mM, KH2PO4 1.25 mM, MgSO4 2 mM, CaCl2 2 mM, NaHCO3 26 mM, glucose 10 mM; pH 7.2, 300 mOsm/l, and positioned on the stage of a vibratome slicer and cut to obtain transverse 350 μm hippocampal slices. Slices were maintained continuously oxygenated for at least 1 h before use. All experiments were carried out at physiological temperature (32 – 34 °C). For experiments, slices were continuously perfused with the solution described above. For experiments, whole-cell patch-clamp recordings were made from pyramidal cells located in the CA1 field of the hippocampus. CA1 pyramidal cells were patched under visual guidance by infrared differential interference contrast microscopy and verified to be pyramidal neurons by their characteristic voltage response to a current step protocol. Neurons were recorded in voltage-clamp configuration with a patch-clamp amplifier (Multiclamp 700B), and data were acquired using *pCLAMP 10.2* software (Molecular Devices). Patch electrodes were pulled from borosilicate glass tubing and had resistances of 4 – 7 MΩ when filled with CsCl 120 mM, HEPES 10 mM, NaCl 8 mM, MgCl2 1 mM, CaCl2, 0.2 mM, EGTA 2 mM and QX-314 20 mM; pH 7.2–7.3, 290 mOsm/l. The firing threshold was measured with *Clampfit* at the potential where the cell fired the first action potential (AP). The data was analyzed using the *Clampfit 10.2* software (Molecular Devices). Resting membrane potential (Vm) was directly measured in current-clamp mode immediately after getting the whole-cell configuration.

### Magnetic resonance spectroscopy (MRS)

Localized [^1^H]MRS was performed at 11.7 tesla using a 117/16 USR Bruker BioSpec system (Bruker BioSpin GmbH, Ettlingen, Germany) interfaced to an advance III console and operating ParaVision 360 V3.6 under TopSpin software (Bruker BioSpin GmbH, Ettlingen, Germany). After fine-tuning and shimming of the system, water signal FWHM values typically in the 15–25-Hz range were achieved. Scanning started with the acquisition of three scout images (one coronal, one transverse and one sagittal) using a 2D-multiplane T2W RARE pulse sequence with Bruker’s default parameters. These images were used to place the spectroscopy voxel of size 1.4 × 1.6 × 1.6 mm^3^ located at the right striatum of the mouse brain or 2.2 × 0.6 × 2.2 mm^3^ located in the cortex (at the midline of the brain), always with care not to include the ventricles in the voxel (the geometry of the voxel was slightly altered to avoid this event, when necessary). At least two ^1^H-MRS spectra were acquired per scanning session per animal (5-month-old animals). The voxel was repositioned, and shimming adjustments were repeated between acquired spectra, when the spectral resolution of the obtained ^1^H spectrum was not good. For ^1^H-MR Spectroscopy, a localized sLASER (semi-Localized by Adiabatic Selective Refocusing) sequence was used. The main acquisition parameters were: echo time (TE) = 18–22 ms, repetition time (TR) = 2,500 ms, and number of averages ranging from 128 to 360 depending on voxel location and acquisition protocol. Spectra were acquired with 2,048 data points, a spectral width of 7,936.5 Hz (≈15.86 ppm), and an acquisition time of 258 ms. Water suppression was applied using a VAPOR module in some acquisitions, while in others no water suppression was used, with a bandwidth of 120 Hz centered at 4.7 ppm. Outer volume suppression (OVS) was enabled to minimize contamination from surrounding tissues. MR spectra were fitted and quantified using LCModel 6.3-1R^50^.

### Matrix-assisted laser desorption/ionization-mass spectrometry imaging (MALDI-MSI)

Male and female control and Gfap-Mpc2-KO mice (N = 8–10 for each condition) were sacrificed and the brain was immediately extracted, divided in half by the hemispheres, embedded in 3 % carboxymethylcellulose (CMC; C4888, Sigma) and fresh-frozen in liquid nitrogen-cold isopentane (277258, Sigma). Blocks of CMC-embedded tissue were sectioned on a *HM 525NX* cryostat (Thermo Fisher Scientific). Sections of 10 μm were mounted onto *MALDI IntelliSlides*™ (1868957, Bruker). One sample of each experimental condition was mounted on the same slide to ensure that all the conditions are represented on a single slide. Sections were immediately put on a desiccator and left in a vacuum environment for at least 30 minutes. Slides were scanned in an *FS120* scanner (Braun) to localize the tissue. They were subsequently prepared for matrix spray on a *HTX M5* Sprayer (HTX Imaging). To prepare the matrix, 4 mg/ml of 1,5-diaminonaphtalene (56451, Sigma) was dissolved in ethanol:H2O:HCl 1 M 9:8:1 (all of them LC-MS grade) and sonicated for 15 minutes. The matrix was taken with a syringe coupled to a 21G needle and it was loaded into the sprayer filtering it with a 0.20 μm polytetrafluoroethylene membrane to ensure that no dust particle or non-dissolved matrix flocculus entered the sprayer. The matrix was applied over the slide at 80 μl/min, 10 psi and 70 °C of nozzle temperature. After the matrix was applied, crystals of phosphorus red were added in a corner of the slide to be used in the next step for the mass spectrometer calibration. Matrix-coated slides were loaded into a *timsTOF fleX MALDI-2* mass spectrometer (Bruker). To perform the MALDI-MSI run, the laser was set at 1 burst of 300 shots with 10,000 Hz frequency. A spatial resolution of 20 μm was used for all the runs. Laser height and position were adjusted for each run, and the phosphorus red was used as a mass calibrant, together with an online calibration provided by the commercial house. The mass spectrometer was set to negative ion mode, detecting m/z values ranging from 50 to 900. Each brain section generated 150,000 individual pixels on average. In every experiment, an area containing only CMC and matrix was analyzed to check for peaks generated by these compounds. Using the *QuPath* v0.5.1 software, hippocampal regions of each brain sample were annotated. Annotations were then exported to the *SCiLS Lab 2024b* v12.01.16074 software (Bruker). Peaks were annotated using a custom library with a mass accuracy of ±10 ppm, and they were quantified without denoising and applying root mean square normalization. Metabolites with high signal in the CMC and matrix area were filtered out, resulting in 113 peaks annotated. For downstream analysis of MALDI-MSI spatial metabolomics data, spatially-resolved root mean square (RMS)-normalized abundance data for all metabolites of interest were first exported as CSV files within *SCiLS Lab* and further analyzed within the R framework (www.R-project.org). RMS-normalized data for each metabolite across each tissue section were first aggregated within the hippocampal subregion previously defined, and annotated accordingly to this region. Aggregation was performed simply by averaging the RMS-normalized data across all pixels in each of these subregions. The subregion-aggregated counts for each metabolite were further log2-transformed, with any zeros first set to 50% of the lowest non-zero value for the corresponding metabolite across all subregions and tissue sections, to avoid numerical overflow errors upon log2-transformation. Statistical analysis was then performed based on a linear categorical model applied on a per-metabolite basis to these log2-transformed, subregion-aggregated counts. To consider the relevance of a detected metabolite, *p-value* together with the biological relevance given the genetic background context, inclusion in a common pathway with a highly significant compound, residing in a similar functional biochemical family with other significant compounds and correlation with other experimental approaches was taken into account.

### [U-^13^C]Glucose tracing using gas chromatography-mass spectrometry (GC-MS)

To assess [U-^13^C]Glucose tracing in cultured astrocytes, a DMEM formulation was prepared consisting of DMEM powder (D5030, Sigma) 8.3 g/l, [U-^13^C6]Glucose (CLM-1396, Cambridge Isotope Laboratories) 5 mM, NaHCO3 26 mM, phenol red (P5530, Sigma) 40 μM; pH 7.4. Astrocytes seeded on 60 cm^2^ plates were washed once with PBS and incubated with DMEM-[U-^13^C]Glucose for 1 h. After incubation, cells were washed with ice-cold PBS. Astrocytes were scraped with 1.5 ml of ice-cold ethanol 70 %. Extracts were centrifuged at 20,000 *g* for 20 min at 4 °C. The ^13^C enrichment of glycolytic products, TCA cycle intermediates and amino acids was measured by GC–MS following a previously described protocol^51^. Lyophilized cell extracts were reconstituted in water, acidified with HCl, and subjected to two ethanol extractions. Metabolites were subsequently partitioned into an organic phase using 96% ethanol/benzene and derivatized with N-tert-butyldimethylsilyl-N-methyltrifluoroacetamide. Samples were analyzed by gas chromatography (Agilent Technologies 7820A, equipped with a J&W HP-5MS column, part no. 19091S-433) coupled to mass spectrometry (Agilent Technologies MSD 5977E). Peak identification and integration from chromatograms were performed using Mass Hunter Workstation software, v B.06.00 (Agilent Technologies), enabling quantification of the detected metabolites. The measured ^13^C enrichment was corrected for natural isotope abundance using unlabeled metabolite standards. Results are expressed as the percentage of labeling of the isotopologue M+X, where M represents the molecular mass of the unlabeled compound and X indicates the number of incorporated ^13^C atoms or as the molecular carbon labelling (MCL), a weighted average of ^13^C-enrichemnt in a given metabolite pool.

### Plasma metabolomics through nuclear magnetic resonance (NMR)

Blood samples from WT and Gfap-Mpc2-KO mice were obtained from the carotid artery. Blood was incubated in ice to let it coagulate and was centrifuged at 1,000 *g* for 5 minutes at 4 °C to separate plasma from red blood cells. Plasma samples were sent to the company *Biosfer Teslab* to perform ^1^H-NMR. Before ^1^H-NMR analysis, 50 μl of murine plasma was added to a standardized 150 μl of human plasma buffer and diluted with 50 μL of deuterated water (D2O) and 300 μL of 50 mM PBS at pH 7.4 into a 5-mm NMR glass tubes. NMR spectra were recorded at 306 K on an *Avance III-600* spectrometer (Bruker BioSciences) operating at a proton frequency of 600.20 MHz (14.1 T). Low molecular weight metabolites were identified and quantified in the 1D Carr-Purcell-Meiboom-Gill spectra using an adaptation of *Dolphin*^52^. Several database engines (*BBioref AMIX* database (Bruker), *Chenomx* and *HMDB*^53^, and literature^54^ were used for 1D-resonances assignment and metabolite identification. After ^1^H-NMR metabolomic characterization, the diluted serum samples were lyophilized and then diluted with 100 μL of 50 mM PBS at pH 7.4 before the lipid extraction using the *BUME*^55^ method with slight modifications. *BUME* was optimized for batch extractions with di-isopropyl ether (DIPE) replacing heptane as the organic solvent. This procedure was performed with a BRAVO liquid handling robot which has the capacity to extract 96 samples at once. The upper lipophilic phase was completely dried in Speedvac until evaporation of organic solvents and frozen at -80 °C until NMR analysis. Lipid extracts were reconstituted in a solution of CDCl3:CD3OD:D2O (16:7:1, vol/vol/vol) containing tetramethylsilane 1.18 mM as a chemical shift reference and transferred into 5-mm NMR glass tubes. ^1^H-NMR spectra were recorded at 286 K operating at a proton frequency of 600.20 MHz using an *Avance III-600* spectrometer. A 90° pulse with water pre-saturation sequence was used. Quantification of lipid signals in ^1^H-NMR spectra was carried out with *LipSpin*^56^, an in-house software based on Matlab. Resonance assignments were done on the basis of literature values^54^.

### Statistical Analysis

For simple comparisons, we used unpaired one-tailed or two-tailed Student’s *t*-test or unpaired Welch’s *t*-test when the variances between populations are different. For other multiple-values comparisons, we used one-way or two-way ANOVA followed by Tukey or Šídák post-hoc test. For the RNA-Sequencing analysis, a Wald test with local fitting was performed. The False Discovery Rate was set at a threshold of FDR<0.05 to consider statistical significance. *p-values* are indicated numerically in the graphs, and all tests used are indicated in each figure legend. . For the determination of metabolic fluxes using radioisotopic tracers, the values of each experimental day were normalized against the control condition. Non-normalized values are shown in the figure legend as mean ± S.E.M. The statistical analysis was performed using the *GraphPad Prism v10.6.0* software. The number of biologically independent culture preparations or animals used per experiment is indicated in the figure legends.

## Reporting summary

Further information on research design is available in the Nature Portfolio Reporting Summary linked to this article.

## Data availability

Source data, statistical table, and uncropped and replicas of the western blots are provided with this paper.

## Acknowledgements

We acknowledge the technical assistance of M. Resch, L. Martin and E. Prieto-Garcia. JPB is funded by the European Research Council (ERC) Advanced Grant NeuroSTARS (101199747), the European Union’s Horizon Europe research and innovation program under the MSCA Doctoral Networks 2021 (101072759; FuEl ThEbRaiN In healtThY aging and age-related diseases, ETERNITY, the Ministerio de Ciencia, Innovación y Universidades/Agencia Estatal de Investigación (MICIU/AEI) (PID2022-138813OB-I00/10.13039/501100011033 and FEDER, UE), Instituto de Salud Carlos III, Plan Nacional de Drogas (2025I002), la Caixa Foundation (grant agreement LCF/PR/HR23/52430016). AAP is funded by the Instituto de Salud Carlos III (PI24/00810, PMP22/00084 and RD24/0009/0005 co-funded by the European Union) FEDER; Junta de Castilla y León (CSI011P23). P.R.-C. is funded by MICIU/AEI (PID2023-152005OB-I00). ARM is funded by the MICIU/AEI (PID2022-136597NB-I00).

## Authors contributions statement

Conceived the idea and designed research: JPB

Performed research: DGR, STS, MAD, LHL, JA, LSO, RL, EF, IMG, ASG, JFG, SPG, IK, MC, MP, BIA, DJB

Analyzed data: JPB, DGR, SE, SJRH, ARM, MP, SMF, PRC, BIA, AA, DJM

Wrote the manuscript: JPB

Edited and approved the manuscript: All co-authors

---

**Extended Data Fig. 1.**
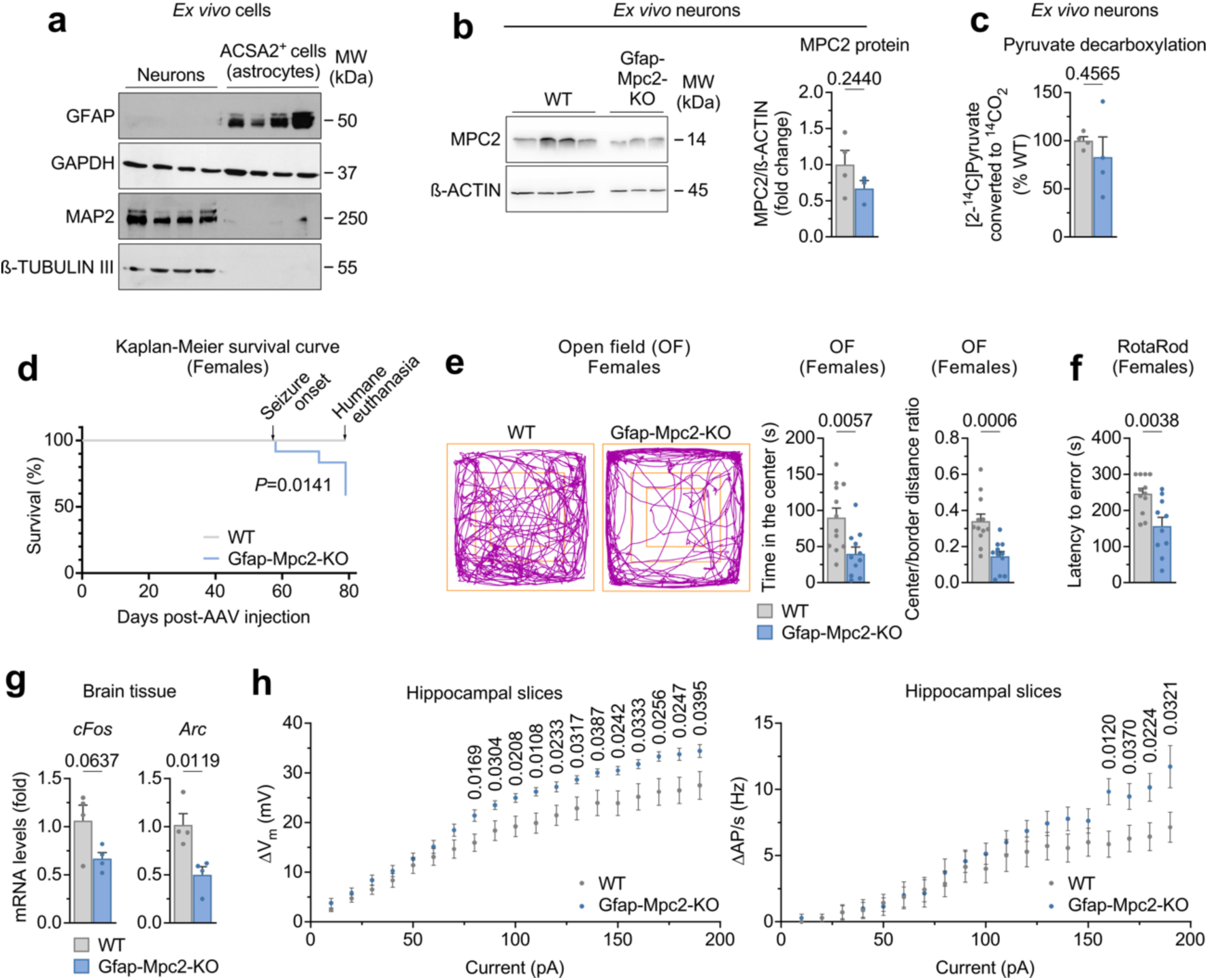
Related to Fig. 1. **(a)** Western blot against GFAP, MAP2 and β-TUBULIN III in immunomagnetically isolated astrocytes and neurons. GFAP was used as an astrocytic marker, MAP2 and β-TUBULIN III were used as neuronal markers and GAPDH was used as a loading control. **(b)** Western blot against MPC2 protein in neurons immunomagnetically isolated from WT and *Gfap-Mpc2-KO* mouse brains; β-Actin was used as loading control (*left*). Quantification of MPC2 protein relative to β-Actin (*right*). Data are mean ± S.E.M. *P* value is indicated; n=4 (WT) or 3 (*Gfap-Mpc2-KO*) mice per genotype; Unpaired Student’s *t*-test, two-tailed. **(c)** Pyruvate decarboxylation flux at the TCA cycle in neurons immunomagnetically isolated from WT and *Gfap-Mpc2-KO*. Data are mean ± S.E.M. *P* value is indicated; n=4 mice per genotype; Unpaired Student’s *t*-test, two-tailed. Non-normalized values are 9.07 ± 1.25 nmol x h^−1^ x mg protein^−1^ for WT and 6.91 ± 1.13 nmol x h^−1^ x mg protein^−1^ for *Gfap-Mpc2-KO* mice. **(d)** Kaplan-Meier survival curve of WT and *Gfap-Mpc2-KO* female mice. *P* value is indicated; n=12 mice per genotype; Logrank test. **(e)** Open field test in WT and *Gfap-Mpc2-KO* female mice. Representative track plots for each genotype are shown (*left*). Quantification of time in the center and center/border distance ratio (*right*). Data are mean ± S.E.M. *P* values are indicated; n=12 (WT) or 11 (*Gfap-Mpc2-KO*) mice; Unpaired Student’s *t*-test, two-tailed. **(f)** Rotarod test in WT and *Gfap-Mpc2-KO* female mice. Data are mean ± S.E.M. *P* value is indicated; n=12 (WT) or 10 (*Gfap-Mpc2-KO*) mice; Unpaired Student’s *t*-test, two-tailed. **(g)** Quantification of mRNA levels of *cFos* and *Arc* genes in brain tissue of WT and *Gfap-Mpc2-KO* mice. Data are mean ± S.E.M. *P* values are indicated; n=4 mice per genotype; Unpaired Student’s *t*-test, two-tailed. **(h)** Electrophysiological recordings in WT and *Gfap-Mpc2-KO* mice neurons from hippocampal slices. Quantifications of the neuron membrane potential (ΔVm; *left*) and action potential frequency (ΔAP/s; *right*) are shown. Data are mean ± S.E.M. *P* values are indicated; n=13 (WT) or 10 (*Gfap-Mpc2-KO*) mice; Unpaired Student’s *t*-test, two-tailed.

**Extended Data Fig. 2.**
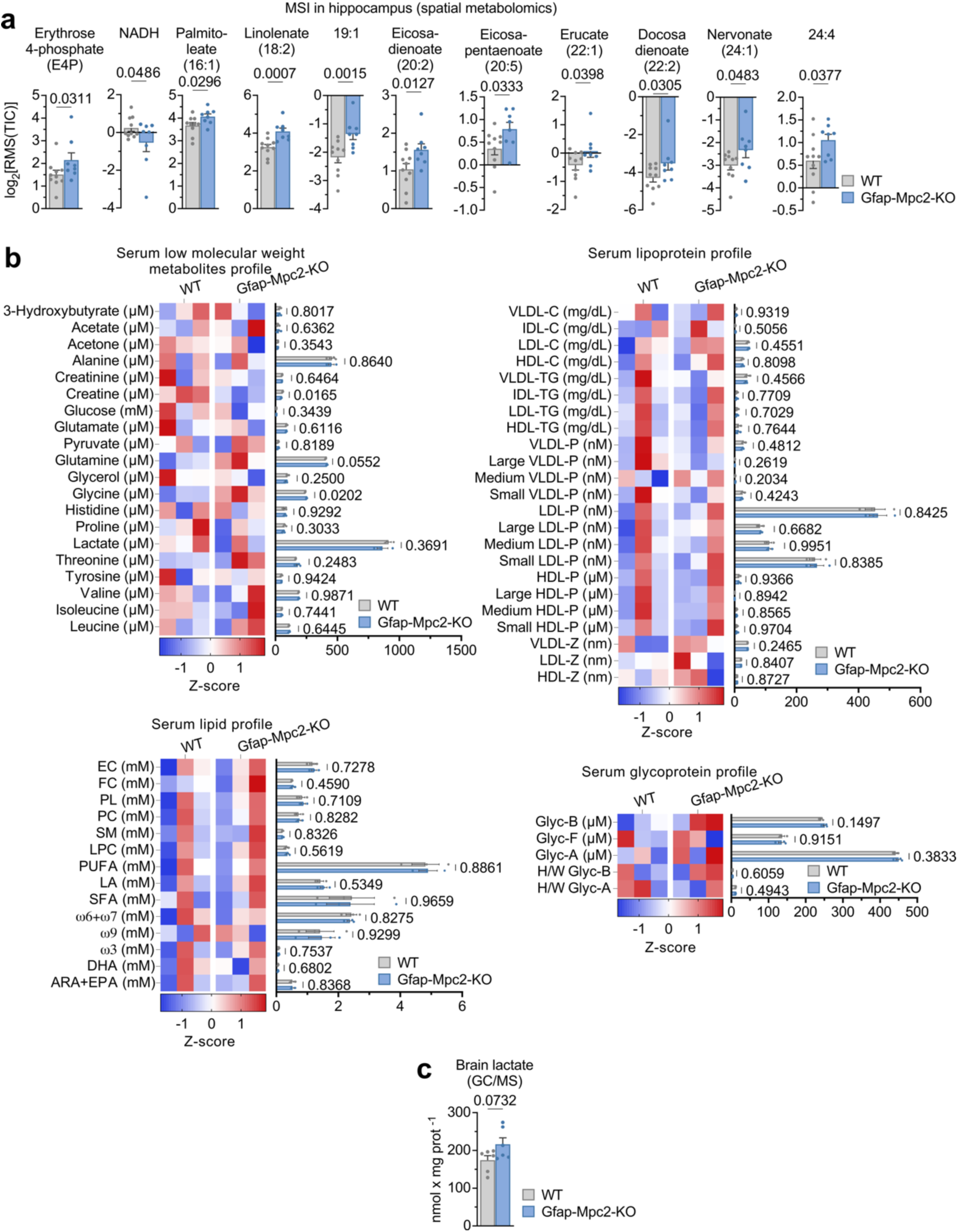
Related to Fig. 2. **(a)** Quantification of selected metabolites detected through mass spectrometry imaging (MSI) in hippocampus of WT and *Gfap-Mpc2-KO* mice. Data are mean ± S.E.M. *P* values are indicated; n=10 (WT) or 8 (*Gfap-Mpc2-KO*) mice; Linear categorical model. (RMS; root mean squared; TIC, total ion count). **(b)** Quantification of metabolites detected through 1H-nuclear magnetic resonance in plasma of WT and *Gfap-Mpc2-KO* mice. Quantifications are shown as heatmaps and absolute values. Data are mean ± S.E.M. *P* values are indicated; n=3 mice per genotype; Unpaired Student’s *t*-test, two-tailed. **(c)** Quantification of lactate levels detected through gas chromatography/mass spectrometry (GC/MS) in whole brain from WT and *Gfap-Mpc2-KO* mice. Data are mean ± S.E.M. *P* value is indicated; n=6 mice per genotype; Unpaired Student’s *t*-test, two-tailed.

**Extended Data Fig. 3.**
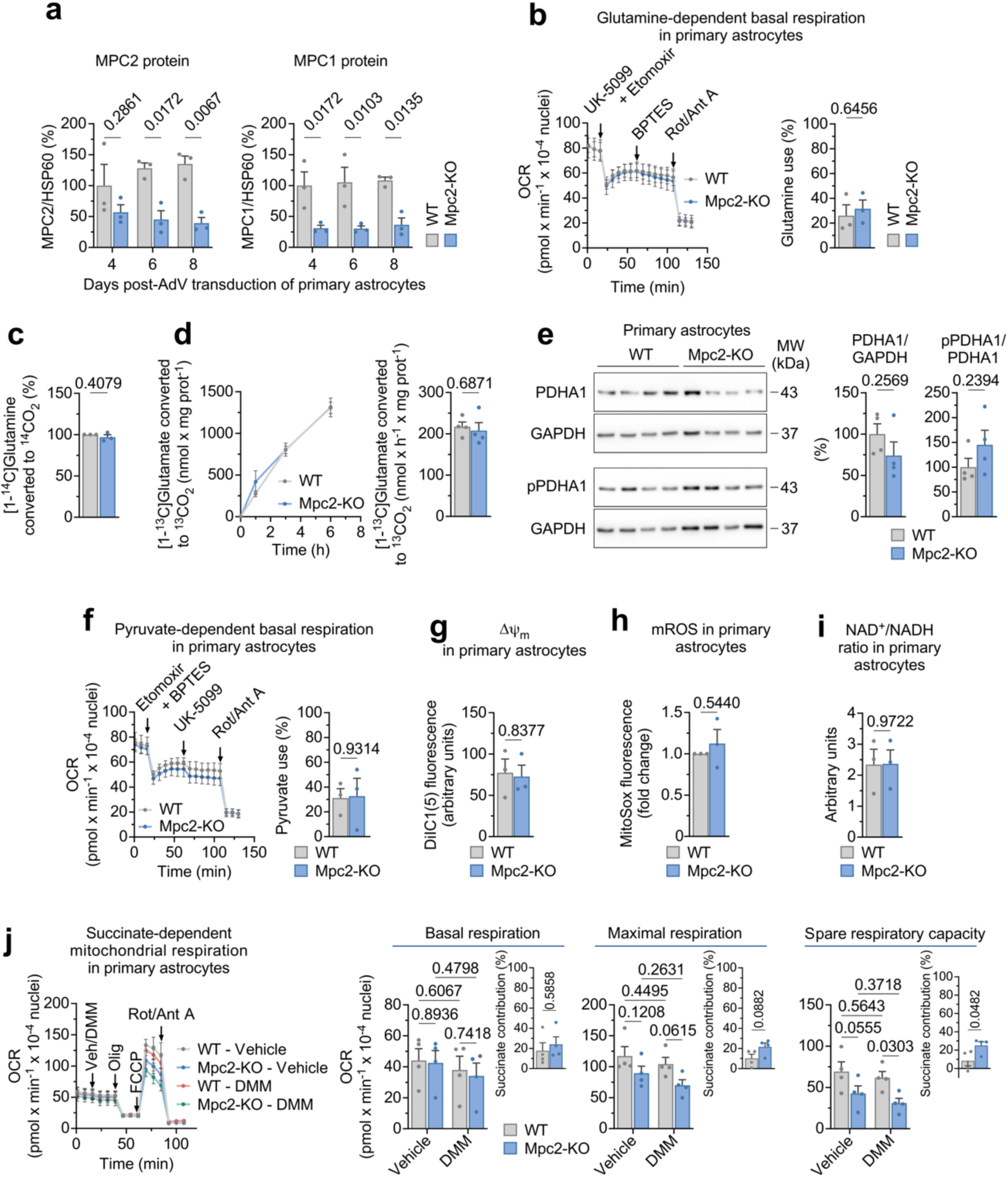
Related to Fig. 3. **(a)** Quantifications of Western blot against MPC2 and MPC1 proteins at 4, 6 and 8 days after AdV-CMV-Cre transduction in immunomagnetically purified mitochondria from primary astrocytes. Data are mean ± S.E.M. *P* values are indicated; n=3 biologically independent samples per genotype; Two-way ANOVA followed by Šidák. **(b)** Glutamine-dependent basal respiration in WT and *Mpc2-KO* primary astrocytes. *Left*: OCR traces; *right*: quantification of glutamine use. Data are mean ± S.E.M. *P* value is indicated; n=3 biologically independent samples per genotype; Unpaired Student’s *t*-test, two-tailed. **(c)** Glutamine decarboxylation flux in WT and *Mpc2-KO* primary astrocytes. Data are mean ± S.E.M. *P* value is indicated; n=3 biologically independent samples per genotype; Unpaired Welch’s *t*-test, two-tailed. Non-normalized values are 185.62 ± 13.44 nmol x h^−^1 x mg protein^−1^ for WT and 179.64 ± 12.21 nmol x h^−1^ x mg protein^−1^ for *Gfap-Mpc2-KO* astrocytes. **(d)** Glutamate decarboxylation flux in WT and *Mpc2-KO* primary astrocytes. *Left:* ^13^CO2 detected levels after 1, 3 and 6 hours of incubation. *Right*: quantification of glutamate decarboxylation flux. Data are mean ± S.E.M. *P* value is indicated; n=4 biologically independent samples per genotype; Unpaired Student’s *t*-test, two-tailed. **(e)** Western blot against PDHA1 and pPDHA1 proteins in WT and *Gfap-Mpc2-KO* primary astrocytes; GAPDH was used as loading control (*left*). Quantification of PDHA1 and pPDHA1 proteins relative to GAPDH (*right*). Data are mean ± S.E.M. *P* values are indicated; n=4 biologically independent samples per genotype; Unpaired Student’s *t*-test, two-tailed. **(f)** Pyruvate-dependent basal respiration in WT and *Mpc2-KO* primary astrocytes. *Left*: OCR traces; *right*: quantification of pyruvate use. Data are mean ± S.E.M. *P* value is indicated; n=3 biologically independent samples per genotype; Unpaired Student’s *t*-test, two-tailed. **(g)** Mitochondrial membrane potential (ΔΨm) levels in WT and *Mpc2-KO* primary astrocytes. Data are mean ± S.E.M. *P* value is indicated; n=3 biologically independent samples per genotype; Unpaired Student’s *t*-test, two-tailed. **(h)** Mitochondrial ROS (mROS) levels in WT and *Mpc2-KO* primary astrocytes. Data are mean ± S.E.M. *P* value is indicated; n=3 biologically independent samples per genotype; Unpaired Welch’s *t*-test, two-tailed. **(i)** NAD^+^/NADH(H^+^) ratio in WT and *Mpc2-KO* primary astrocytes. Data are mean ± S.E.M. *P* value is indicated; n=3 biologically independent samples per genotype; Unpaired Student’s *t*-test, two-tailed. **(j)** Succinate-dependent mitochondrial respiration in WT and *Mpc2-KO* primary astrocytes. *Left*: OCR traces; *right*: quantification of basal respiration, maximal respiration and spare respiratory capacity; *right, insets*: succinate contribution to each metabolic parameter. Data are mean ± S.E.M. *P* values are indicated; n=4 biologically independent samples per genotype; Two-way ANOVA followed by Tukey or Unpaired Student’s *t*-test, two-tailed.

**Extended Data Fig. 4.**
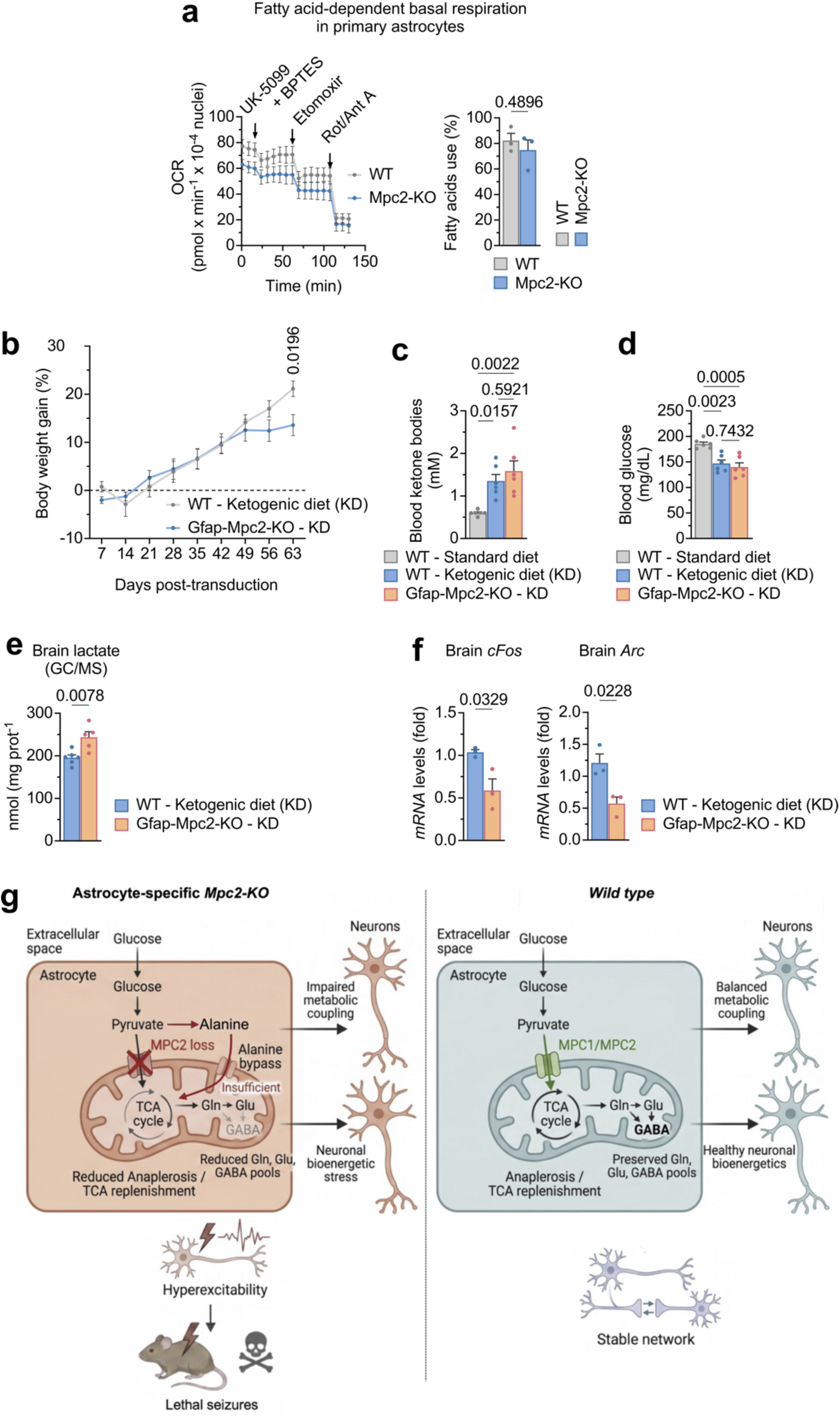
Related to Fig. 4. **(a)** Fatty acid-dependent basal respiration in WT and *Mpc2-KO* primary astrocytes. *Left*: OCR traces; *right*: quantification of fatty acids use. Data are mean ± S.E.M. *P* value is indicated; n=3 biologically independent samples per genotype; Unpaired Student’s *t*-test, two-tailed. **(b)** Body weight gain of WT and *Gfap-Mpc2-KO* mice undergoing a ketogenic diet. Data are mean ± S.E.M. *P* value is indicated; n=6 mice per genotype; Unpaired Student’s *t*-test, two-tailed. **(c)** Blood ketone bodies in WT and *Gfap-Mpc2-KO* mice undergoing a ketogenic diet. Same-age same-sex mice under a standard diet were used as basal measurement. Data are mean ± S.E.M. *P* value is indicated; n=6 mice per genotype; One-way ANOVA followed by Tukey. **(d)** Blood glucose in WT and *Gfap-Mpc2-KO* mice undergoing a ketogenic diet. Same-age same-sex mice under a standard diet were used as basal measurement. Data are mean ± S.E.M. *P* value is indicated; n=6 mice per genotype; One-way ANOVA followed by Tukey. **(e)** Quantification of lactate levels through gas chromatography/mass spectrometry (GC/MS) in whole brain from WT and *Gfap-Mpc2-KO* mice undergoing a ketogenic diet. Data are mean ± S.E.M. *P* value is indicated; n=6 (WT) or 5 (*Gfap-Mpc2-KO*) mice; Unpaired Student’s *t*-test, two-tailed. **(f)** Quantification of mRNA levels of *cFos* and *Arc* genes in brain tissue of WT and *Gfap-Mpc2-KO* mice undergoing a ketogenic. Data are mean ± S.E.M. *P* values are indicated; n=3 mice per genotype; Unpaired Student’s *t*-test, two-tailed. **(g)** Proposed schematic representation summarizing the phenotypical alterations occurring in the *Gfap-Mpc2-KO* mice.

